# One Stop MS Shop (OSMSS): statistical analysis and interpretation of quantitative proteomics data for non-coders

**DOI:** 10.64898/2026.11.13.766418

**Authors:** Erez Cohen, Pierre A Coulombe

**Affiliations:** Department of Cell & Developmental Biology, University of Michigan Medical School, Ann Arbor, MI; Department of Dermatology, University of Michigan Medical School, Ann Arbor, MI; Rogel Cancer Center, University of Michigan, Ann Arbor, MI

**Keywords:** Quantitative proteomics, differential abundance analysis, R Shiny, Graphical User Interface, limma, proDA, SAINTexpress, TMT, DIA, DDA, phosphoproteomics

## Abstract

Open-source statistical methods for proteomic differential abundance analysis are well established. Normalization, missing-value handling, and the statistical model must suit the experimental design and change how proteins are called. Together with the frequent need for data reformatting, coding and bioinformatics expertise, these elements limit the bench scientist’s analysis of datasets. Here, we present One Stop MS Shop (OSMSS), a no-code R Shiny application that parses Proteome Discoverer, MaxQuant, FragPipe, DIA-NN or Spectronaut search results for user-driven interactive exploration through transformation and differential analysis. Upon import, OSMSS records the experimental design through user questions, selects the analysis workflow, identifies normalization requirements, runs established statistical methods, and records every analytical step in a case-specific methods text. OSMSS supports label-free data-dependent and data-independent acquisition (DDA, DIA) and tandem mass tag (TMT) group comparisons, affinity-purification comparisons, and DIA and TMT phosphosite analyses. OSMSS runs on local devices or servers and provides volcano plots, heatmaps, trend analysis, kinase prediction, pathway enrichment and kinase activity inference. In seven benchmarks, OSMSS performed comparably to matched reference workflows. OSMSS lets bench scientists apply established statistical models and associated tools transparently and without coding, enabling a more informed analysis of their data.

## Introduction

Quantitative proteomics studies often ask which proteins or their modifications differ in abundance between conditions. As such, multiple methods for quantifying differential abundance are established and freely available as R packages^1,2^. They include limma, developed for gene expression and applied to proteomics, proDA, which models missing values^3^, SAINT used for analysis of affinity purification–mass spectrometry (AP-MS) datasets^4^, and MSstats for abundance comparisons in label-free and label-based data^5^. Across these applications, data transformation, normalization and formatting, which make samples comparable, must suit the dataset and foster reliability^6^. The statistical approach must also suit the acquisition method and replicate number^7^. Additionally, missing values arise from different underlying factors^8^ and the fraction of missing values differs between DDA, DIA and multi-batch TMT data^9–11^. For example, a protein missing from every sample in a group may occur at a level below the instrument’s limit of detection^12^. Imputation methods, if used to replace missing values with estimates, each assume one of these factors^8^. Choices at any of the above steps can change which proteins (or modifications) are called differentially abundant^13^.

Choosing the right methodology, transformations and data formatting is thus crucial for robust analysis of mass spectrometry (MS) data. However, this process requires users to understand intricacies of both statistical approaches and available software. To illustrate, in group comparisons of many proteins, moderated tests such as limma, which shrink each protein’s variance toward a pooled estimate^2^, are more powerful and stable than ordinary t-tests^14^. Affinity purification illustrates how strongly experimental design matters to method selection: nonspecific binders are commonly filtered with SAINT, which scores candidates interacting with a bait against negative-control purifications^4^. In such a case, true interactors are expected to be absent from controls and have no values in the control samples^15^. Thus, a test of group means would not call them significant without imputation of these missing values^15^. These methodological differences extend to how such samples should be normalized, as SAINTexpress scores spectral counts or intensities directly^16^. Such methodological differences exist across the broad spectrum of proteomics, including analysis of tandem mass tag (TMT) single and multi-plex experiments, analysis of data-independent acquisition (DIA), or post-translational modifications (PTMs)^17,18^. Such a multitude of tools and methods is a testament to how the fields of proteomics and biostatistics have grown and, together, have greatly expanded our ability to explore and analyze proteomics data over the past two decades.

For researchers outside these fields, however, these choices present a challenge and a steep learning curve: finding the correct analysis method and software, reformatting search results for it, choosing parameters, and interpreting outcomes without clearly stated methodology. PRIDE, described by its authors as the world’s leading proteomics repository, now receives over 500 datasets a month on average^19^. Analysis of any of these datasets requires reporting each analytical step, which can be tedious, error prone and skill intensive^20,21^. Processing is often delegated to specialists^22^. For non-experts, published guides explain how to calculate fold changes and P values in a spreadsheet^22^. Multiple labs have thus taken to develop graphical tools for differential analysis, including LFQ-Analyst^23^, MSstatsShiny^24^, Phospho-Analyst^25^, FragPipe-Analyst^15^, Perseus^26^, amica^27^, ProteoArk^28^, ProStaR^29^, and MaxQuant and MSstats in Galaxy^30^ (Table 1)^31–38^. Several of these interfaces guide the user through the analysis: PMart blocks methods unsuited to the data type^39^, and Eatomics turns a hypothesis into a model formula^40^. Alternatively, others record the analysis itself such as MSstatsShiny^24^, Perseus^26^, MS-DAP^41^, PMart^39^ and Galaxy^30^, and several produce reports that list the selected settings, as in FragPipe-Analyst^15^, amica^27^, Eatomics^40^ and prolfquapp^42^. Of note, the recent combination of apps in the Analyst Suites of the Monash Proteomics and Metabolomics Platform^43^ provides a large number of tools to analyze input across data types (LFQ-, Phospho-, FragPipe-, TMT-, DIA- and Pathway-Analyst). These suites provide a powerful tool for user analysis and a variety of normalization and imputation methods^44,45^, as well as standardized limma differential abundance analysis^15,23,46^. As such, they provide a step forward towards accessibility for non-expert users. However, despite their accessibility, users are still left with a large number of choices that require their explicit understanding of biostatistics, including imputation type or normalization type, as well as the formatting and input methods. For example, a user uploading the outcome of a single plex TMT experiment might not know the meaning of “internal reference scaling (IRS) normalization” preselected by TMT-Analyst, and whether or not it is relevant to their data^44^. Thus, while graphical interface tools have become available over the past decade, a remaining gap for bench scientists is guidance for issues such as the selection of appropriate methodology, across experiment types, using native search-engine exports and, most importantly, receiving a transparent, case-specific record of how their data were processed and analyzed, including the checks and choices made, in explanatory text rather than code.

**Table 1.** Features of OSMSS and existing tools for downstream analysis of quantitative proteomics data. The table compares features, not performance. Cells are based on publications, documentation and the software versions current in November 2026. The Analyst Suites column covers the six apps of the Monash Proteomics and Metabolomics Platform (LFQ-, Phospho-, FragPipe-, TMT-, DIA- and Pathway-Analyst) [15, 23, 25, 43–46]. Abbreviations: DDA, data-dependent acquisition; DE, differential expression; DIA, data- independent acquisition; FC, fold change; FDR, false discovery rate; GSEA, gene set enrichment analysis; IMD, independence of missing data; IRS, internal reference scaling; KSEA, kinase–substrate enrichment analysis; ORA, over-representation analysis; PCA, principal component analysis; PTM, post-translational modification; QC, quality control; SASA, solvent-accessible surface area; TMT, tandem mass tag. Footnotes: a, available in the current software version but not described in the cited paper; b, cells refer to the MaxQuant, MSstats and MSstatsTMT tools, and the Galaxy platform hosts further tools, for example for phosphopeptide analysis; c, includes add-on packages built on prolfqua: prolfquapp, prolfquasaint (SAINTexpress) and prophosqua (PTM; no peer-reviewed publication); d, no graphical interface, so the cell describes the generated report.

Here, we present One Stop MS Shop (OSMSS), a no-code browser application built with R Shiny to give non-expert molecular scientists this guidance and allow users a single graphical approach for a large variety of open-source statistical analysis software. OSMSS reads native search results from Proteome Discoverer, MaxQuant, FragPipe, DIA-NN and Spectronaut (including exports through Perseus). Upload wizards ask about the experimental design and from user-supplied answers, OSMSS chooses an analysis route, normalizes (if the export is not already normalized), and runs an established statistical model. OSMSS covers four kinds of experiments: (1) label-free group comparisons from data-dependent acquisition (DDA) or DIA; (2) TMT comparisons within one plex or across several; (3) affinity purification of a bait against negative controls; and (4) DIA and TMT experiments on phosphosites.

OSMSS also writes a rule-based methods text from the settings that ran and links the results to interpretation tools including gene set enrichment analysis and kinase activity inference (PhosR)^47^, all within the application. It also offers kinase prediction tools (The Kinase Library^48,49^) that require no user data and allow hypothesis generation. OSMSS combines established statistical models, including limma, with normalization and handling of missing values guided by the data type and experimental design. To validate OSMSS’s approach, we compared it with reference workflows, established methods run on the same data, in seven benchmarks with known answers (Table 2). Under these tests, OSMSS was within or above the reference range on most measures. Last, OSMSS provides an easy import/export function that allows users to reanalyze and share their data and, importantly, record the analysis pipeline, all within a locally saved small .zip file that holds an R data (RDS) file. To increase accessibility, transparency and reproducibility of published results, users can also upload this .zip with their dataset to PRIDE, a member of the ProteomeXchange consortium^19,50^. This allows data to be reanalyzed in OSMSS to generate hypotheses or replicate analysis by other labs rapidly. We thus propose OSMSS as a user interface for researchers that provides guidance, statistical analysis, and an interface to explore both private and publicly available data from search files. OSMSS is available as a free repository on GitHub in both an R Shiny server version and a locally run application.

**Table 2.** The seven benchmarks: data, input, datasets and comparisons, reference workflows and main call rule. Dataset identifiers and sources are in the Experimental Section (Benchmark Datasets) and in Supplemental Data 1 (Datasets). Except in the knockout, which tested normalization itself, the reference workflows started from OSMSS’s input file and normalized it the same way. Abbreviations: DDA, data-dependent acquisition; IRS, internal reference scaling; PTM, post-translational modification; ROTS, reproducibility-optimized statistical testing; SAM, significance analysis of microarrays.

## Experimental Section

### Implementation and Architecture

OSMSS 3.1 is a Shiny application developed under R 4.6.1^51^ and shiny 1.14.0^52^. The interface uses bslib 0.12.0 for theming^53^, plotly 4.12.1 for graphics^54^ and DT 0.34.0 for tables^55^. Data are held in QFeatures 1.22.0^56^ and SummarizedExperiment 1.42.0^57^ objects. Summarization uses MsCoreUtils 1.24.0^58^, and imputation uses imputeLCMD 2.1^59^ and impute 1.86.0^60^. Structures are read with bio3d 2.4-5^61^ and displayed with NGLVieweR 1.4.0^62^, an R interface to the NGL viewer^63^. These versions, and limma 3.68.5^1^, are the ones used here, for development and for the reference workflows. OSMSS’s benchmark results were generated under R 4.1.2 and limma 3.50.1, the versions on our server (Supporting Information Text). OSMSS runs with the packages installed on the user’s computer, and the generated methods text records the R version and the installed versions of limma, proDA and QFeatures.

OSMSS code has six layers (Figure S1), drawn from its call graph. Upload wizards parse each export, record the experimental design and write reactive objects, which are then read by the analysis, interpretation, output and methods-text modules. Analyses can be saved and reopened under Saved Experiments as a .zip file. OSMSS runs on a server or any computer with R (across Mac, Windows and Linux operating systems); more than 8 GB of memory is recommended.

### Input

Currently, OSMSS upload wizards read native exports of Proteome Discoverer^64^ (protein- level data or nested protein-peptide export), MaxQuant^65^, FragPipe^66^ (with MSFragger^67^ for peptide identification and IonQuant^68^ for label-free quantification), DIA-NN^69^ and Spectronaut^70^. For MaxQuant label-free data, OSMSS reads the raw protein-group Intensity in proteinGroups.txt, which MaxQuant does not normalize. It uses LFQ intensity, MaxQuant’s label-free quantity normalized across runs^71^, only when no raw Intensity columns exist, and the methods text then reports so. Reverse decoy hits and groups identified only by site are removed. OSMSS detects from evidence.txt whether match-between-runs, which transfers peptide identifications between runs, was enabled, and reports it without changing the analysis. The TMT wizard accepts Proteome Discoverer protein or nested protein–peptide exports, or a flat peptide table from other software. The Label Free PTM wizard accepts only Spectronaut’s default precursor report plus a protein-level export. The TMT PTM wizard accepts Proteome Discoverer peptide-group files, one phosphopeptide-enriched and one global file per plex to allow adjustment of each phosphosite for its parent protein’s abundance.

For DDA, one upload box recognizes the file from its column headers. It accepts Proteome Discoverer protein exports, MaxQuant proteinGroups.txt (followed by evidence.txt) and FragPipe combined_protein.tsv. FragPipe input also requires its fragpipe.workflow file. Similar to the MaxQuant process, OSMSS reads the per-sample Intensity columns from FragPipe combined_protein.tsv and removes decoy (“rev_”) rows. From fragpipe.workflow, OSMSS reads the FragPipe version, whether IonQuant normalization was on or not, match-between-runs and the peptides used. A FragPipe file is refused when IonQuant was not run or when labeling was used.

For DIA, one upload box recognizes four inputs: (1) the Spectronaut Pivot Report; (2) a Pivot Report exported through Perseus^26^; (3) DIA-NN’s protein-group matrix (report.pg_matrix.tsv); and (4) FragPipe DIA output, which is DIA-NN’s matrix produced within FragPipe. OSMSS recognizes a Perseus export by the type prefixes Perseus adds to column names. DIA-NN’s matrix is already filtered at a 1% protein-group false discovery rate. Spectronaut run-level (long) reports, which are typically larger than a pre-set 1 GB upload limit, are refused, and the user is directed to the Pivot Report or the OSMSS_Global.rs custom Spectronaut export scheme. DIA-NN’s main report and precursor matrix are refused with directions to the protein-group matrix. Label-free DDA files are directed to the DDA upload.

### Analysis Routes

OSMSS normalization and statistical model selection follow a pre-set of rules depending on experimental design and data type. Search files already normalized are not normalized again. OSMSS supports statistical analysis and data exploration of (1) Label-free DDA data; (2) AP-MS quantitation; (3) DIA protein intensities; (4) TMT single and multi-plex intensities; and (5) phospho-proteomics analysis using absolute PTM and protein-adjusted statistics.

### Analysis Route: label-free DDA

The label-free DDA route is adapted from proDA and follows the workflow in its documentation, which log2-transforms the intensities and normalizes them by the median before the model is fitted^72^. Label-free DDA data from MaxQuant (Intensity) or Proteome Discoverer (Abundance) are always normalized by the global median or an invariant-protein panel based on experimental design and context. FragPipe Intensity is not normalized again when fragpipe.workflow shows that IonQuant normalization was applied (“on” by default^73^), unless the wizard selects the invariant-protein panel. When only a Proteome Discoverer peptide file is supplied, peptides are robustly summarized (using MsCoreUtils) to proteins before normalization. The two normalizations, global median and invariant-protein panel, suit different designs. The global median aligns the median of all proteins across runs. The wizard selects it when the user reports quantitative differences only, as in drug treatments or time courses. When the user reports that proteins are expected to be entirely absent in one condition, as in knockout, knockdown or gene deletion experiments (but not classic AP-MS), the wizard selects the invariant-protein panel instead. The invariant-protein panel holds proteins detected in every run, highly abundant and stable within conditions, and differing little between conditions once any offset shared by the bulk of proteins is removed. If too few proteins qualify for the panel, OSMSS falls back to the global median (thresholds in Supporting Information Text). The Confirm & Summarize step tells the user which normalization will be applied. Proteins with fewer than two observations in either of the compared conditions are kept but flagged as LowEvidence, which does not change the fit. For differential abundance, comparisons are then tested with proDA 1.26.0, which models missing values with a probabilistic dropout model instead of imputing them^3,72^. P values are BH-adjusted (Benjamini–Hochberg method)^74^, and results are reported at adjusted P alone, without a fold-change filter.

### Analysis Route: affinity purification

For affinity purification, the label-free wizard offers several routes depending on the user’s answer regarding the experimental design. The wizard asks what the control sample contains. A non-specific capture control, processed by the same protocol without the bait (for example IgG, or the tag alone), leads to SAINTexpress^16^, briefly described in the next paragraph. A control from a different cellular fraction (for nuclear, chromatin or organelle pulldowns), or the same bait pulled down in another biological condition (to compare interactomes across conditions), leads to proDA with invariant-protein panel normalization.

For affinity purification, the label-free wizard offers SAINTexpress 3.6.3^16^, which scores the bait against negative controls^4^ without normalization by OSMSS, because SAINTexpress normalizes intensities internally^16^. OSMSS parses the upload and writes SAINTexpress’s three input files (bait.dat, prey.dat and interaction.dat). The bait file marks each purification as bait or control, the prey file lists the detected proteins, and the interaction file holds each protein’s intensity in each purification sample (each bait or control purification). OSMSS then runs the compiled SAINTexpress program, which the user installs from its source (Data and Software Availability), in the background. The results table reports each protein’s SaintScore and Bayesian false discovery rate from SAINTexpress, ranked by SaintScore. Thresholds, input scaling and fallbacks to proDA are given in the Supporting Information Text.

SAINTexpress models a single bait-versus-control comparison, but a design with a non-specific control may include more conditions. In such a case, the wizard relates the need to select only 2 conditions (bait and control) in order to use SAINTexpress. proDA is offered when users, receiving this alert, still require comparing multiple conditions across affinity purifications.

### Analysis Route: DIA global protein changes

The DIA protein route is adapted from limma’s empirical Bayes test^1^, as used previously in published proteomics workflows^75^. Protein-group quantities from Spectronaut (pre-normalized by default) are log2-transformed and not normalized a second time. To check whether the export is already normalized, OSMSS takes the proteins quantified in every run and compares each run’s median log2 intensity with those of the other runs under the same condition. A spread (standard deviation) of 0.02 log2 or less is consistent with normalized data, and more than 0.03 suggests the data were not normalized. OSMSS shows this result next to a checkbox, ticked by default, stating that the data were normalized in Spectronaut. The check guides the user’s choice, which matters most for data processed with older Spectronaut versions (before version 15^76^). Starting with version 15, Spectronaut’s default cross-run normalization changed from global normalization to an automatic choice that uses local normalization for fewer than 500 runs^76^. If the user wishes to override the default by unticking the checkbox beside the run-median check, OSMSS median-centers the log2 values once. For DIA-NN, the search file protein-group matrix is read directly from the DIA-NN report (report.pg_matrix.tsv) or from FragPipe DIA output and can be uploaded with DIA-NN’s report.log.txt or FragPipe’s fragpipe.workflow. Either format records whether DIA-NN normalized across runs (current default unless run with --no-norm). OSMSS log2-transforms the DIA-NN matrix and median-centers it only when DIA-NN did not normalize. Without a log file, OSMSS assumes DIA-NN’s default, and the user confirms it with the checkbox located beside the run-median check on the user menu. In the DIA protein route, a protein is tested if it has been quantified in at least 50% of samples in both conditions compared. Due to the nature of missing values in DIA, only proteins that pass this detection filter and are missing at most one value per condition are imputed with k-nearest-neighbor imputation^60,77^. Other proteins that pass the filter, with more than one missing value in a condition, are fitted with limma^1^ on observed values and are not filled otherwise. Proteins that pass the filter are tested per limma recommendations, with empirical Bayes moderation, intensity-dependent variance trend^1,75^ and robust hyperparameter estimation^78^, which limits the effect of proteins with unusually variable measurements. Proteins entirely failing the detection filter (<50% of samples in either condition) are tested with Fisher’s exact presence/absence test, which queries whether detection differs between the conditions, with BH adjustment within that set^74^. These proteins appear in the “not missing at random” (NMAR, often termed MNAR) analysis tab, with its own tables and download. All conditions share one model unless their typical within-condition standard deviations differ more than 1.5-fold; then only the two compared conditions are fitted. P values from the limma fit are adjusted and reported. For DIA-NN input, the methods text also reports the DIA-NN version, the matrix used, whether the normalization setting came from the log file or from the user, and whether DIA-NN ran within FragPipe.

### Analysis Route: TMT single and multi-plex global protein changes

The TMT protein route is also adapted from limma’s empirical Bayes test^1^. TMT single plex reporter intensities are log2-transformed and median-centered once, after verification that data have not already been normalized as described in the DIA route. In multi-plex data, the bridge design applies median centering, followed by IRS on the mean of each plex’s pooled reference channels if a reference channel exists^79^. For multi-plex experiments with no pooled channel, a balanced design provides an alternative analysis and, within this design, applies median centering only. In this design, OSMSS checks that every plex holds the same number of samples of each condition (or rejects the input). In single plex data, k-nearest-neighbor imputation^60^ fills values only for proteins missing at most one value in every condition. The balanced design uses k-nearest-neighbor imputation within each plex, and the bridge design is not imputed. A protein is tested only if it is quantified in at least 50% of plexes (multi-plex data) and in at least 50% of the replicates (channels) of at least one condition; others are listed with their coverage and the reason in a Not Tested & Low Coverage table provided to the user. The same tab lists tested proteins that are quantified in at least half of the channels of only one of the two compared conditions. These proteins are flagged as low coverage in the results, and those with no value at all in one compared condition receive no P value. All conditions are fitted in one limma model with sample weights. MSstatsTMT also fits all conditions together after its reference-channel normalization^80^. In the balanced design, the plex enters the model as a covariate, a block effect for plex-to-plex differences, as in DEqMS^2^ and msqrob2TMT^81^. Empirical Bayes moderation uses the variance trend and robust estimation, as in the DIA route.

### Analysis Route: PTM, TMT and DIA (absolute and adjusted)

Phosphosites are tested with limma, as for proteins^1^, and the Adjusted view described below relates each phosphosite to its parent protein, adopting the pipeline used in msqrob2PTM analysis^82^. In analyzing PTM data, users input both a phospho-enriched and a global (protein-level) file from the same samples. The analysis routes map each peptide to a phosphosite, where the decision of which residue contains the event relies on the search software’s localization probability, the confidence that the modification occurs on that residue, with at least 75% site confidence used as the default. This generates a phosphosite matrix used in the following steps. In the TMT PTM route, the phosphosite matrix is median-centered on peptide-level medians. This matrix is centered separately from the global protein matrix, which comes from the second user input file and is centered on its own column medians. Multi-plex data then use the IRS bridge or the plex covariate, similar to protein analysis. TMT sites are not imputed, and a site needs at least two values in both compared conditions to pass the filter.

In the Spectronaut DIA PTM route, median polish, a summary that resists outliers, combines each site’s precursors into one value per sample^58^. The PTM and global exports, which Spectronaut normalizes across runs by default, are not normalized again. A run-median check is shown for each export, and if the user unticks its box, OSMSS median-centers that matrix and the adjusted values. For a phosphosite to be compared in downstream analyses, the summarized peptides containing the site need values in at least half of a user-defined condition’s samples, with a minimum of two. k-nearest-neighbor imputation is applied only to phosphosites missing at most one value per condition^60^. In both TMT and DIA PTM routes, phosphosites that fail the filter are tested for presence versus absence by Fisher’s exact test on their detection counts in the Absolute view, BH-adjusted within that set.

OSMSS generates three views for each PTM comparison. The Absolute view tests the site’s own abundance. The Protein view tests the parent proteins of the global file. The Adjusted view subtracts, in each sample, the parent protein’s log2 intensity in the matched global run from the site’s, re-centers the result (unless both Spectronaut exports were already normalized) and fits limma^82^. On the log2 scale, subtracting the protein’s value gives the phosphosite’s ratio to its protein – it becomes normalized to its parent protein^82^. In limma, applying such an adjustment requires the analysis to preserve robust estimation but remove the variance trend parameter. In contrast, the Absolute and Protein fits use the variance trend and robust estimation parameters. P values are adjusted and reported as above for each of the different views. The Supporting Information Text gives each route’s settings.

### Generated Methods Text

Each analysis writes its settings and counts to a provenance record, which fixed sentence templates turn into a user-friendly methods section. The text covers the input, the features removed, match-between-runs status, normalization and its check, filters, numbers of untested and imputed values, the model and its empirical Bayes settings, and multiple testing.

### Tools Beyond MS Analysis

Gene set enrichment analysis^83^ uses clusterProfiler 4.20.0^84^ on MSigDB gene sets^85^ through msigdbr 26.1.1^86^. Trend clusters use fuzzy c-means soft clustering (e1071 1.7.17^87^), with the fuzzifier estimated by the method of Schwämmle and Jensen^88^, as in Mfuzz^89^. Coverage maps draw UniProt annotations^90^ with drawProteins 1.32.0^91^. Kinase-Site Analysis infers kinase activity from the Absolute or Adjusted PTM results with the kinase–substrate scoring of PhosR, implemented in OSMSS with PhosR 1.22.0 reference data^47^, including per-sample activity scores. The PTM differential expression view (PTM DE in the menu) exports the results in a format for kinase-substrate enrichment analysis^92,93^.

The prediction tools run on a protein or residue and need no MS data. Kinase Prediction scores sites against The Kinase Library’s Ser/Thr and Tyr specificity matrices^48,49^ (v1.2.0; Apr 15, 2025, in this study) and overlays known sites from PhosphoSitePlus^94^ (2022 release, obtained January 2026, in this study). Each user downloads these files from their source (Data and Software Availability), so the versions in use may differ. Motif and binding predictions query Scansite^95^ and 14-3-3-Pred^96^. Structures originate from the AlphaFold Protein Structure Database^97,98^. Solvent accessibility is computed on the AlphaFold structure, read with bio3d^61^, by the Shrake–Rupley method^99^. The conservation track aligns orthologs from the Ensembl REST service^100^ with Clustal Omega^101^ through msa 1.44.1^102^.

The PTM database holds sites built from FragPipe psm.tsv files with their peptide.tsv^66,67^, one row per observed modification, with experimental metadata entered by the user. The user keeps it as an exported, dated zip file and loads it again by upload. The Results section describes how this database can be used for hypothesis generation based on preliminary PTM data across a large number of experiments. These experiments are the user’s own FragPipe searches, of enriched or unenriched samples. Their modified sites are added over time to the one locally kept database, where a site seen repeatedly across experiments can then be found.

### Benchmark Datasets

All benchmark datasets that we used to test OSMSS performance are public. DDA, DIA and TMT single plex datasets come from OpDEA, a published set of spike-in benchmarks with known changes^13^. The DDA benchmark used 12 datasets with 22 comparisons, read from MaxQuant raw Intensity. Four are human–yeast–E. coli hybrid proteomes from PXD028735^103^, and one is a timsTOF hybrid proteome from PXD014777^104^. Three are yeast backgrounds spiked with UPS1, a standard mixture of 48 human proteins^105^: PDC000006 from the Proteomic Data Commons^106^, PXD002099^107^ and PXD001819^108^. Two are human–yeast mixtures (PXD007683^109^; PXD036134^41^) and two are human–E. coli mixtures (PXD018408^7^; PXD021425^10^).

The benchmark exercise for DIA used 6 datasets with 14 comparisons, quantified with Spectronaut 18 in directDIA mode^70^. These are one each from PXD028735^103^, PXD018408^7^, PXD019777^10^ and PXD036134^41^, and two UPS1-in-E. coli dilution series from PXD026600^105^.

The benchmark exercise for TMT single plex used 5 datasets with 15 comparisons: a human–E. coli TMT10 mixture (PXD013277^2^), a human–yeast TMT11 mixture (PXD007683^109^) and three acquisitions of the HYpro16 two-proteome standard^110^. For these datasets and for the phospho TMT benchmark, the MaxQuant output was converted to the Proteome Discoverer format that the OSMSS TMT wizards read (Supporting Information Text). The TMT multi-plex benchmark used the MSstatsTMT SpikeIn-5mix data (MassIVE.quant RMSV000000265 and RMSV000000266), UPS1 proteins spiked into a HeLa background^80^. These data were acquired in MS2 and SPS-MS3 modes, which are two ways of measuring TMT reporter ions. The data hold five TMT10 mixtures (plexes). Each mixture held the four UPS1 amounts twice each and a pooled reference sample twice, and each mixture was run three times in each mode. We treated one run of all five mixtures as one experiment, as in a five-plex study, which gave six datasets (two modes × three runs). In each dataset, every UPS1 amount was compared with every other, for 36 comparisons in total.

The benchmark exercise for semi-synthetic knockout (for testing invariant protein normalization) simulated known shifts into six timsTOF HeLa runs^10^ with 4,540 eligible proteins. The runs were split into control and knockout groups, and a known subset of proteins was shifted in the knockout runs only. Proteins that shifted below the detection limit became missing. The four scenarios used the same six runs and differed only in the share of eligible proteins that was shifted: none, 26%, 46% or 66%. In each scenario with changed proteins, most shifted proteins decreased in the knockout runs. Shifting many proteins changed each knockout run’s total protein. To simulate equal loading, each knockout run was then rescaled to the control runs’ total protein. This rescaling moved every protein in the run, including those that did not change. Unchanged proteins were thus left with loading offsets of +0.09, +0.30 and +0.59 log2 (higher in the knockout runs), in order of increasing share changed.

The phospho TMT benchmark used PXD007145, in which yeast phosphopeptides were diluted 1:4:10 into a HeLa background^111^. Each of its 17 TMT10 runs was one dataset, with 51 comparisons in total. The phospho DIA benchmark used PXD014525^112^, one dataset with 10 comparisons. The authors’ Spectronaut project was reprocessed in Spectronaut 21.1^70^ without the HeLa-only and yeast-only runs, with no imputation, local cross-run normalization and a q-value cutoff of 0.01. Spectronaut’s localization cutoff was set to 0, so OSMSS’s 75% threshold was the only localization filter. Neither phosphosite study measured an unenriched proteome, so the global file required by the wizard was a stand-in built from the existing data (Supporting Information Text). Both phosphosite benchmarks, phospho TMT and phospho DIA, were scored in the Absolute PTM view.

### Benchmark Reference Workflows

The reference workflows started from OSMSS’s input file and, except in the knockout, normalized it the same way (Table 2). In DDA they were limma^1^, proDA^3^, DEqMS^2^, Student’s t-test and ROTS^113^; DIA and both TMT protein benchmarks used these workflows without DEqMS. The phosphosite benchmarks used the same four plus MSstatsPTM 2.14.0^18^, and phospho DIA also included significance analysis of microarrays (SAM), a permutation test, through samr 3.0.1^114,115^. MSstatsTMT 2.20.0^80^, run with its default settings on the same peptide-level input, was the only reference workflow for OSMSS’s TMT peptide routes. On the multi-plex data, the MSstatsTMT 2.20.0 reference agreed with the authors’ deposited MSstatsTMT 1.6.2 results^80^. The limma reference used plain empirical Bayes without trend or robust estimation. Last, because the simulated knockout data test normalization, its references were limma, DEqMS, proDA, the t-test and ROTS with or without median normalization, plus proDA variants, limma after left-censored imputation^59^ and DEP^116^. In the scenario with 66% of proteins changed, DEP was not run, so 15 reference workflows remained. SAM was excluded from the phospho TMT ranges because, with three replicates per condition, its smallest adjusted P value was 0.10 in every run.

### Scoring and Statistics

For benchmark tests, true changes were assessed as previously done^13^, at either their published mixing ratios or spike-in amounts. A feature was called at adjusted P ≤ 0.05 alone (DDA, invariant-protein knockout, DIA and TMT multi-plex), or at adjusted P ≤ 0.05 and a fold change of at least 1.5-fold (TMT single plex, phospho DIA and phospho TMT benchmarks). In some benchmarks, normalization shifts the unchanged background in every workflow, and without the fold-change filter many unchanged proteins or phosphosites reach significance, so that about half or more of every workflow’s calls were false. In those benchmarks (TMT single plex, phospho DIA and phospho TMT), the fold-change filter is the main rule. The filter also mirrors how users read a volcano plot, by fold change as well as by P value.

Each call was a true or a false positive. The false-discovery proportion (FDP) is the share of calls that are false, and sensitivity is the share of true changes called. In addition to FDP and sensitivity, the normalized Matthews correlation coefficient (nMCC), (MCC + 1)/2 on a 0–1 scale, combines all four outcomes of calling at adjusted P 0.05 into one score^117^. The partial area under the curve (pAUC), with McClish standardization^118^, measures how well the P values rank true changes (above unchanged features) at false-positive rates up to 0.01 and 0.05. Fold-change correlation represents the Spearman rank correlation between estimated and true fold changes. Using these metrics, OSMSS’s value was compared against the reference values, as means across datasets or within one dataset. OSMSS’s performance was then ranked as above, within or below the reference. Counts are given as a percentage of the median reference (the median of the reference workflows’ values), and rates and scores as percentage points (pp). OSMSS was compared with the median reference by two-tailed paired t-tests over datasets, with Holm–Šidák adjustment across each benchmark’s eight metrics^119^. Statistical analysis was done using GraphPad Prism v11. Per-dataset values for OSMSS and every reference workflow are in Supplemental Data 1.

## Results and Discussion

### OSMSS guides users from search results to interpretation

Across the analytical workflows and upon receiving an uploaded search result file, OSMSS judges five elements towards robust statistical analysis (Figure 1A). These are (1) the input column indicating which quantity to read; (2) normalization of the data; (3) handling of missing values; (4) decision on the statistical model to fit; and (5) which rule calls a protein changed. Each of the first four choices can change which proteins are called^7,13,41^. To guide users through these processes, OSMSS turns these choices into questions about experimental design. Background scripts also parse the search file directly, reading its column names, descriptions and verifying whether the uploaded data are pre-normalized. OSMSS uses this information to suggest an analysis route suited to the experimental design.

**Figure 1.**
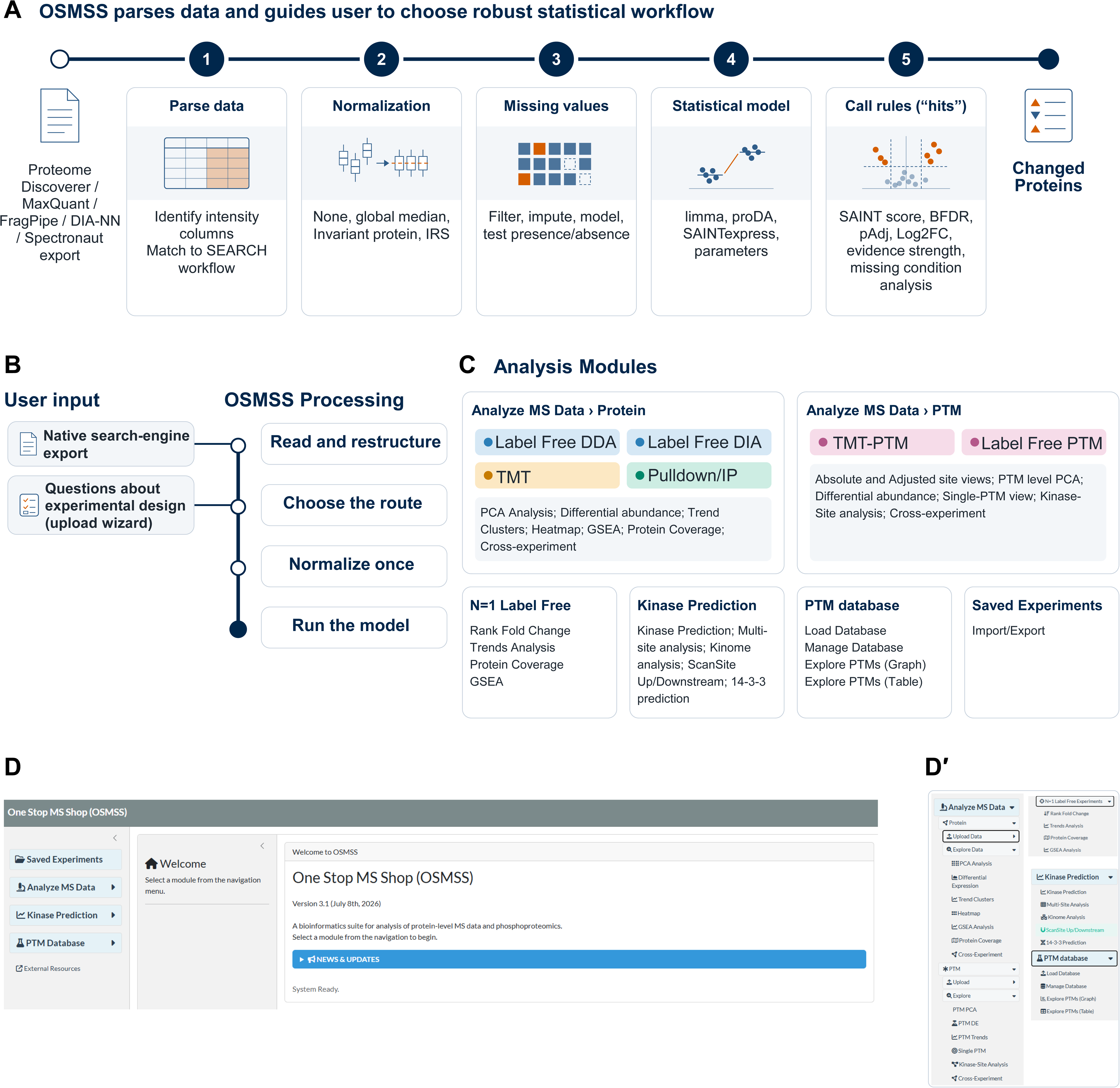
From search results to interpreted results. (A) The five decisions that follow a search engine’s export (Proteome Discoverer, MaxQuant, FragPipe, DIA-NN or Spectronaut). (B) User-provided native search-engine export and design Q&A in the upload wizard. OSMSS then reads and restructures the data, chooses the route, normalizes once and runs the model. (C) The analysis modules and tools provided in the OSMSS menu. (D) Screenshot of the OSMSS welcome screen. (D′) The expanded navigation menu.

To initiate data analysis, users upload a native export of their search results (Figure 1B; *Experimental Section*). Due to the large number of available native exports from Spectronaut, we have generated two export-ready files for .sne reports (OSMSS_Global.rs, OSMSS_PTM.rs), provided with OSMSS, which allow streamlined export of pivot Spectronaut files to OSMSS. Users are instructed to upload a file according to the experiment type (DDA, DIA, TMT, DIA-PTM or TMT-PTM). The label-free DDA and DIA uploads recognize the file and prompt users for unresolved information.

The menu groups the uploads and analyses into three analysis modules (Figure 1C,D,D′): (1) Protein analysis, with uploads for Label Free DDA, Label Free DIA and TMT data; (2) PTM analysis, with uploads for TMT-PTM and Label Free PTM (DIA) data; and (3) N=1 Label Free Experiments, which supports rank fold change, trends, protein coverage and gene set enrichment analysis for unreplicated preliminary DDA data analysis run via Proteome Discoverer. Together, the modules cover four kinds of experiments: label-free group comparisons acquired by DDA or DIA; TMT group comparisons within one plex or across several; affinity purification of a bait against negative controls; DIA and TMT experiments that quantify phosphosites. Additional modules provide users with options for prediction-based analysis and data import, including (4) Kinase predictions modules, (5) a cross-experiment dataset to store preliminary PTMs from phospho- and non-phospho-enriched experiments for hypothesis generation and (6) an import/export module which allows re-analysis of datasets, and ease of data sharing within labs and across collaborations.

### The experimental design determines the analysis route

Once a user uploads their search data, the upload wizard queries the user about the following elements: (1) sample to group and condition allocations; (2) experimental design (e.g. pulldown or immunoprecipitation); and (3) information regarding run design (e.g. whether a TMT plex carries pooled reference channels). From the answers, OSMSS reads and restructures the export, chooses the analysis route, normalizes the data once (or not at all if pre-normalization has occurred or is detected) and runs an established statistical model (Figure 1B). However, a new version of a search program can change its default settings, such as normalization. To guide users on whether their data might have already been normalized, OSMSS checks the median of each run, tells the user whether the data look normalized, and offers a checkbox to override the default if necessary. The wizard explains to the user that normalizing pre-normalized data a second time, with a different method, can shift proteins that did not change. For DIA-NN exports without a log file, the user confirms the normalization setting through a checkbox, similar to the approach for Spectronaut exports (Figure 2A).

**Figure 2.**
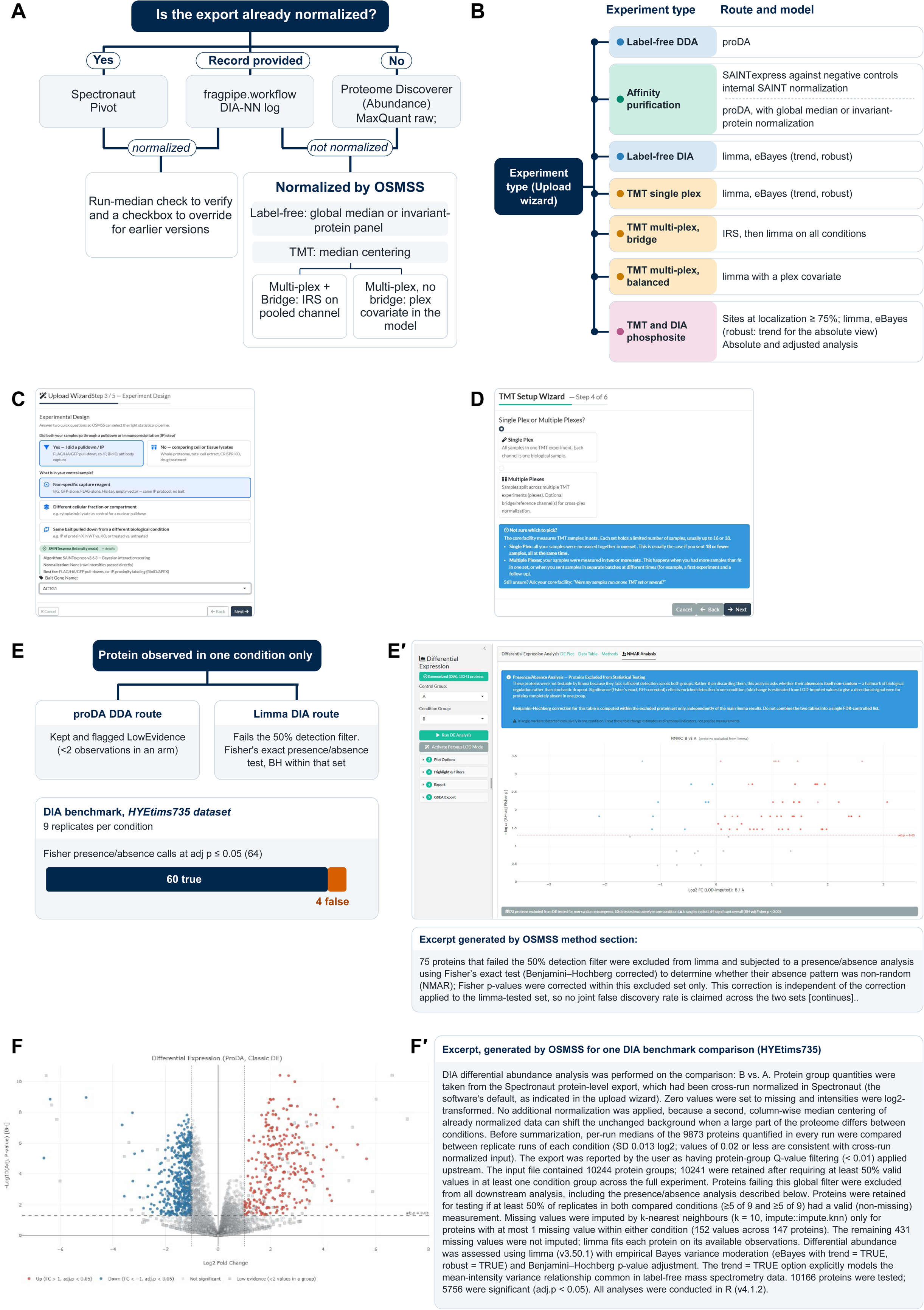
How the experimental design determines the analysis. (A) Normalization decision: whether the export is already normalized, and what OSMSS does in each case. (B) Analysis route and statistical model for each experiment type. (C) Upload wizard example, design step for a pulldown or immunoprecipitation with a non-specific control. (D) TMT setup wizard. (E) Proteins observed in one condition only, in the proDA (DDA) and limma (DIA) routes. Inset: true and false Fisher presence/absence calls at adjusted P ≤ 0.05 in the HYEtims735 DIA benchmark dataset (9 replicates per condition). (E′) NMAR Analysis tab, with its methods-text excerpt; triangles mark proteins detected in one condition only. (F) Volcano plot of the HYEtims735 DDA benchmark dataset (MaxQuant; condition B vs condition A, 7 replicates each): log2 fold change against −log10 of the Benjamini–Hochberg (BH)-adjusted P value. Red (up) and blue (down) points pass both dashed thresholds (adjusted P < 0.05 and |log2 fold change| > 1); light grey circles, not significant; grey squares, LowEvidence proteins (fewer than two values in a condition), drawn grey whatever their fold change and P value. (F′) Generated methods-text excerpt for one DIA benchmark comparison (HYEtims735).

Global median normalization is a standard step in published workflows^2,12,72^. In addition to the global median normalization routes, OSMSS provides an invariant-protein panel option. This panel considers that global median normalization assumes most proteins do not change between conditions^41^. However, when many proteins change in the same direction (for example, after a knockout), the median itself moves. Aligning the medians then shifts the proteins that did not change. Consequently, the invariant-protein panel is a context-specific alternative to the global median, where a set of stable proteins selected by OSMSS from the data is used for normalization. Normalizing on a stable reference set has a precedent in the stably phosphorylated sites used by PhosR^47^ and in Normics, which normalizes using a subset of proteins that it selects from the data as unchanged^120^. The panel aims to keep proteins that did not change at a log2 fold change near zero, even when many other proteins shift in one direction. The normalization then does not create false changes in these proteins. We further tested this using the knockout benchmark below, in which the more proteins changed in one direction, the larger the panel’s advantage over global median normalization.

After data transformation and normalization, each experiment type then goes to a published model (Figure 2B), and the Experimental Section describes the process used for each route. Label-free DDA data are tested with proDA. Alternatively, when the user chooses affinity purification^4,16,121^ as their design (Figure 2C), SAINTexpress gives the user a score for each prey against their own control purifications. SAINT analysis has been offered through ProHits^122^ and the CRAPome website^121^ and through the REPRINT website^121^. Each resource takes the data in its own input format, such as separate bait, prey and interaction files in SAINT ^123^. OSMSS does the formatting for the user (see Experimental Section). The user then runs the statistical analysis and graphs directly from the native search export.

In contrast to DDA, DIA protein data are tested with limma^1,2,14^. Two choices in the DIA route shape the limma statistical analysis. First, to let the pooled variance estimate follow a protein’s intensity, OSMSS keeps the variance trend^75^. The authors of limma describe the trend as important because many gene expression technologies give less reliable data at low intensities or abundances^1^. Second, OSMSS reports every tested protein at adjusted P alone and leaves any fold-change filter to the user. To bring forward the larger changes users typically look for, the volcano plot colors the proteins above a 2-fold change threshold. The cost of these choices, extra calls with small fold changes, is measured in the benchmarking effort related below.

For TMT, users are first guided to determine whether their TMT data are single- or multi-plex (Figure 2D). In multi-plex experiments, the wizard-issued query regarding pooled reference channels sets the design. In the bridge design, each plex carries pooled reference channels. To put the plexes on one scale, IRS scales each plex to these channels^11,79^. In TMT multi-plex experiments, where no pooled IRS reference exists (“balanced design”), OSMSS adds the plex as a covariate in the limma model (see Experimental Section).

In the PTM routes, peptides are mapped to sites, filtered through the localization filter and tested with limma, as shown in Figure 2B. Filtering on it lets the user rely on the position of each tested site. As in the DIA and TMT protein routes, sites that fail the detection filter are not dropped; a presence/absence test checks them instead (Experimental Section). The Adjusted view tests whether a given phosphosite changes beyond its parent protein^17,18,82^, so that the user can tell a change in phosphorylation status from a change in the amount of protein. Since after the adjustment, each site’s value is centered near zero, a limma variance trend has no intensity to follow, and the Adjusted view is fitted without it. OSMSS’s adjustment follows the order of msqrob2PTM^82^ and the approach of Wu et al., who calibrated phosphopeptide ratios by protein levels^17^. Other published tools also adjust for the parent protein and may use different methodologies to do so, as this remains an evolving field^15,18,25,46^. Adjusting a phosphosite for its protein relies on protein-level data from the same samples, as in Phospho-Analyst^25^. PTM analysis in general is also limited by the low abundance of modified proteoforms and by the few observed peptides that span each modification site^18^. Within one peptide, OSMSS keeps both residues of a doubly phosphorylated peptide (multiplicity 2, M2), as assigned by the search software. However, phosphosites on different peptides, which are common when peptides are short, cannot be linked, and whether they occur on the same protein molecule (cis) or on different molecules (trans) cannot be determined^124^.

### Proteins observed in only one condition remain transparent to the user

Proteins observed in only one condition need their own handling. A mean-based test cannot call them without imputation^15^, and a linear model cannot be fitted when a group has no observation^12^. OSMSS therefore handles these proteins explicitly but maintains transparency to the user instead of dropping them, as these proteins could be important for hypotheses and follow-up experiments (Figure 2E). In proDA routes, OSMSS keeps such proteins in the differential abundance plots yet flags them as “LowEvidence” for the user. In the DIA global protein route, proteins that do not pass the detection filter are tested for presence versus absence, with Benjamini–Hochberg correction^74^, and appear in the NMAR analysis tab (see Experimental Section). This transparency, and the ability to detect these proteins, matter to the user. A protein present in samples of interest (e.g. lesion) but absent from the control (e.g. non-lesion) might be excluded from the model, yet it may be the protein that matters most for the user’s hypothesis. The NMAR analysis lists and tests it instead of dropping it (Figure 2E,E′). In the DIA benchmarks, this test made 69 calls in the two datasets with 9 and 8 replicates per condition, and 64 of them were true (64 calls, 60 true, in HYEtims735; Figure 2E). With 3 replicates per condition, the test cannot reach significance. However, even in smaller sample sizes, the NMAR analysis still lists these proteins and their test. This listing provides users information otherwise dropped or hidden. As with the DIA NMAR table, the TMT “Not Tested & Low Coverage” table (Experimental Section) allows users to see proteins that otherwise would not appear in the volcano plot and its estimates. This allows users to fully explore their dataset, and no protein is dropped without being counted in the generated methods text.

### Every analysis is recorded in a generated methods text

Once data upload has been completed, users are prompted to either export their transformed, OSMSS-ready summarized data for future import and sharing (“Saved Experiments”) and/or explore their results through several modules. These output modules include (a) principal component analysis (PCA), which includes coefficient of variation and screen plot analysis (principal components vs eigenvalues); (b) differential expression (volcano plot, tables and NMAR analysis; Figure 2F); (c) heatmaps (z-score or absolute values); (d) trend cluster analysis for multi-condition experiments; and (e) single PTM analysis for absolute/adjusted PTM analysis. Across all modules, a methods text is generated from the analysis, relating how the data were processed from upload to user output.

As noted in the Introduction, reporting every step of a data analysis is tedious and error prone, and a fraction of published studies leave normalization, imputation or statistical parameters absent or vague^20,21^. The generated methods text addresses this incomplete reporting by recording each analysis step. Unlike reports that list the selected settings, the text is specific to each dataset: it gives the counts at each filter, the result of the normalization check, and the reason for each choice or fallback. Figure 2F′ shows an excerpt for one DIA benchmark comparison. As a user-first interface, OSMSS prioritizes recording the methods text as prose, similar to that used in a manuscript methods section. This allows users to read, check, and adapt the processing of their data for a manuscript without the need to fully comprehend the underlying code. This also provides users, peer reviewers, and readers with context to understand how the data were processed without a significant amount of field expertise. As such, users do not need to rely on overly brief statements and instead can provide a detailed account of how their data were analyzed.

### Tools for exploration and interpretation of results

Across the output types provided to users, a large number of visualization options are integrated in submenus via the Plotly R-package. These include the ability to highlight proteins (or PTMs) by gene names or accessions, on-mouse hover information, filtering by data (such as the number of peptide-spectrum matches, log2 fold change, presence/absence or protein group) and exports as CSV.

Several additional analysis tools are provided to complement differential abundance (Figure 3A,B). PCA relates whether samples coalesce by condition, and how the analysis changes when applying batch correction or the IRS bridge (uncorrected PCA view). The interactive heatmap shows proteins as row z-scores or as abundance values, for each replicate or averaged by condition. To help the user find proteins that rise or fall together in experiments with several conditions, trend clusters group proteins by their profile across conditions^89^. Gene set enrichment analysis^83–86^ tests whether the changes concentrate in known gene sets such as pathways. Protein coverage maps show which parts of a protein are covered by the captured and analyzed peptides, with UniProt-defined domains overlaid^90^ (Figure 3B; data from PXD082237^125^), and a cross-experiment view compares results between analyses. This view takes in the standard differential expression output from OSMSS across all analysis routes (for example, 3 different SAINTexpress runs across bait/controls) and generates an interactive Venn diagram allowing multi-experiment exploration.

**Figure 3.**
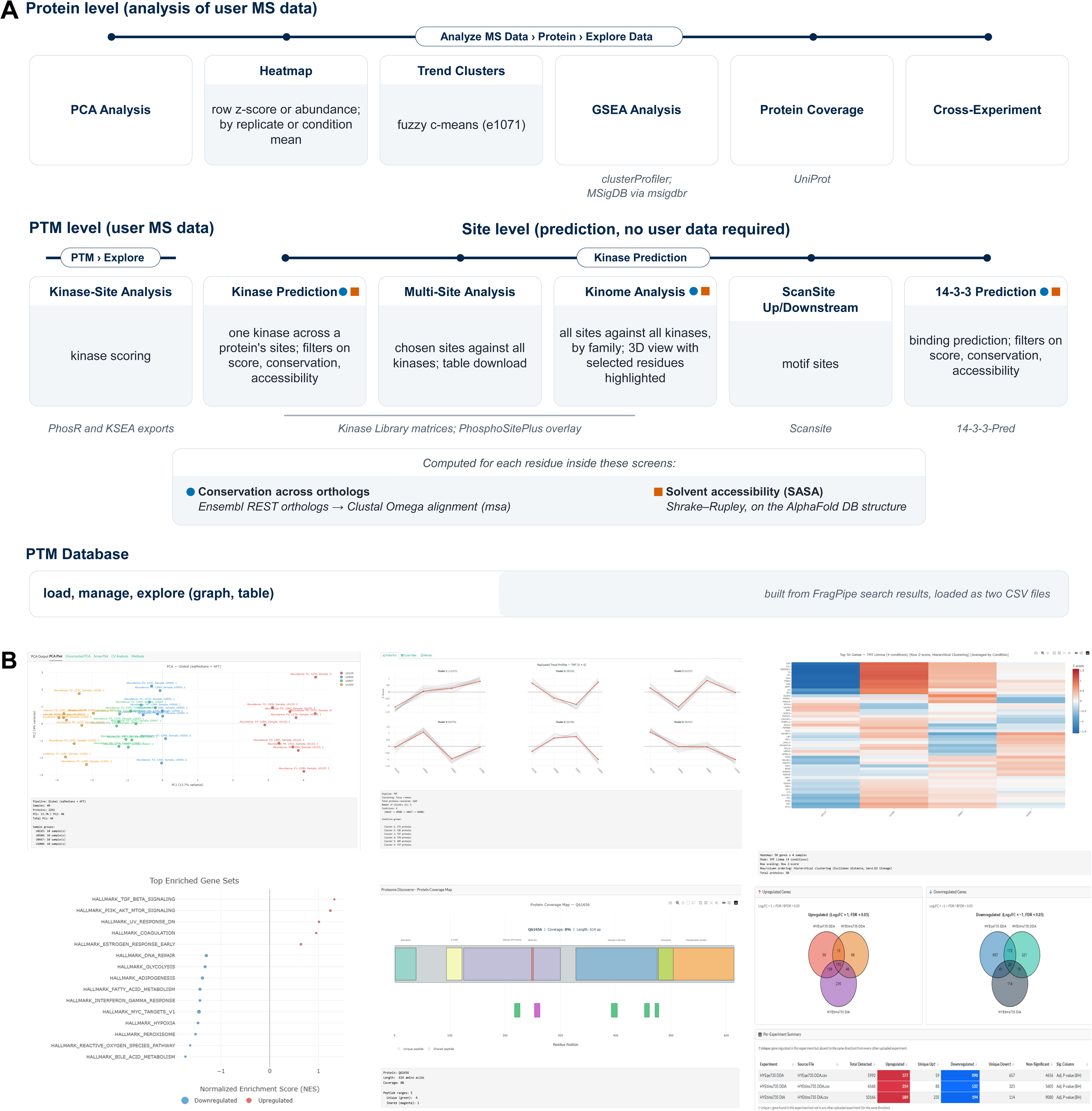
Tools for interpreting differential results and hypothesis generation. (A) Protein-level tools, which use the user’s MS data (Analyze MS Data>Protein>Explore Data); site-level tools, prediction based and requiring no user MS data (Kinase Prediction); PTM-level tools, which use the user’s phosphosite results (PTM › Explore: Kinase-Site Analysis and the PhosR and KSEA exports); and the PTM Database. The resources each tool uses are named in the panel: clusterProfiler [84], MSigDB [85] through msigdbr [86], e1071 [87] with the Mfuzz fuzzifier estimate [88, 89], UniProt [90], The Kinase Library [48, 49], PhosphoSitePlus [94], Scansite [95], 14-3-3-Pred [96], Ensembl [100], AlphaFold DB [98] and PhosR [47]. KSEA [92, 93] is not run within OSMSS. (B) Screenshots of the six protein-level tools: PCA (after plex batch correction), trend clusters and heatmap, run on the MSstatsTMT SpikeIn-5mix TMT multi-plex benchmark data [80]; GSEA overview (top MSigDB Hallmark gene sets by normalized enrichment score), on the B vs A result of the HYEtims735 DIA dataset [103]; protein coverage, from the Proteome Discoverer export of PXD082237 [125]; and the cross-experiment Venn diagram of the B vs A results from three acquisitions of the same human– yeast–E. coli samples (HYEtims735 DDA, HYEqe735 DDA and HYEtims735 DIA [103]).

For phosphosite experiments, Kinase-Site Analysis in the PTM menu takes the user’s own Absolute or Adjusted PTM results and infers kinase activity from the whole differential result with the kinase–substrate scoring of PhosR^47^. This also allows users to view kinase enrichment score per sample, for example predicted kinase enrichment per patient across multiple diseases. The PTM results can be exported in a format for kinase-substrate enrichment analysis^92,93^. None of these tools was benchmarked here and each has been developed separately and published before, namely gene set enrichment analysis^83^ with clusterProfiler^84^, fuzzy c-means clustering as in Mfuzz^89^, PhosR^47^ and kinase-substrate enrichment analysis^92,93^. OSMSS integrates these tools together into a single interface, adds layers to allow users to explore and output the results of these tools through user-interactive plots and filters, and by doing so improves users’ ability to integrate and explore their data across several established tools.

### Additional tools requiring no MS data: kinase prediction and PTM exploration

To assist hypothesis generation and data exploration through kinase prediction tools, the ‘Kinase Prediction’ menu gathers resources that do not require a user-uploaded dataset. The user enters a protein’s UniProt ID and gene symbol and, where needed, the residue numbers of the sites of interest. A phosphosite can come from the user’s own PTM results or from any other source or hypothesis. The kinase scores originate from The Kinase Library’s peptide specificity matrices for Ser/Thr and Tyr kinases, which describe the sequences each kinase prefers^48,49^. Three views answer different questions, as follows: (1) The Kinase Prediction view, which shows which S/T/Y sites in a given protein are predicted to be acted upon by a kinase of interest. This tool further allows users to check a prediction against sites already reported via PhosphoSitePlus^94^ -it provides scores per residue from The Kinase Library (TKL), or TKL and PhosphoSitePlus overlaid; (2) Multi-Site Analysis, which scores selected phosphosites against all kinases using The Kinase Library matrices; and (3) an exhaustive per-protein Kinome Analysis, which scores all S/T/Y sites against all kinases, grouped by kinase family, for a whole-kinome view of one protein. This view further provides a 3D interactive image of the protein as modeled by AlphaFold^97,98^, where the user can further identify which residue is of interest, and highlight it on the protein structure to better understand where their PTM of interest might place on the protein.

Within these modules, OSMSS calculates two scores for each candidate phosphosite in addition to the kinase scores. The two scores help the user choose candidate sites of interest to follow up, as structural and evolutionary information are used together to prioritize phosphosites^126^. The first, a structural score, is the residue’s solvent accessibility, computed on the protein’s 3D structure from the AlphaFold Protein Structure Database^97,98^. Phosphorylation sites tend to be more solvent-accessible than other serines, threonines and tyrosines^127^. The second, an evolutionary score, is the site’s conservation across orthologs. Conserved phosphosites are, on average, more likely to have functional significance^128,129^. In the Kinase Prediction view and in 14-3-3 Prediction, described below, the user can show only phosphosites above a chosen solvent accessibility or conservation. In Kinome Analysis, only conservation can be used this way.

Two further prediction resources sit beside the kinase tools and tell the user, in one application, what else may recognize the same phosphosite. Scansite Up/Downstream queries Scansite at scansite4.mit.edu, which identifies short motifs around the site that modular signaling domains recognize or that protein Ser/Thr or Tyr kinases phosphorylate^95^. 14-3-3 Prediction predicts whether 14-3-3 proteins may bind the phosphosite^96^.

Last, OSMSS provides a “PTM database”, a locally saved database which stores preliminary observations rather than predictions. In contrast to the prediction tools above, the database holds PTM sites actually observed in the user’s own experiments. Such observations need attention, as modified proteoforms are of low abundance^18^, and PTMs found in experiments that were not phospho-enriched may not be well quantified. Individually, such PTMs may potentially be false positives^130^. However, how often a site is identified across samples is informative about its potential function^126^. The database takes in FragPipe psm.tsv files with their peptide.tsv files from the user’s own searches, after FragPipe’s false discovery rate filtering. As the user adds experiments over time (phospho-enriched or not, and of varying origin), the database tracks these PTMs across experiments for hypothesis generation. To check whether a site of interest was seen in other experiments, users can load, manage and explore the database as a cross-experiment graph or a table.

### Benchmarking OSMSS routes against matched reference workflows

Across datasets, OSMSS did not perform significantly worse than the median reference workflow in any benchmark, and in DDA and phospho TMT it performed significantly better on several metrics. When compared dataset by dataset individually, 430 of 496 results (87%) were within or above the reference range, and 124 (25%) were better than every reference workflow. Importantly, the benchmarks test whether the guidance, analytical routing and interface provided by OSMSS cost statistical robustness, not whether OSMSS does better than utilizing packages directly through code. OSMSS was scored in seven benchmarks with known answers (Table 2). Five were at the protein level: label-free DDA, an invariant-protein knockout, DIA, TMT single plex and TMT multi-plex. The remaining two were at the phosphosite level: phospho TMT and phospho DIA. In each benchmark except the knockout (which tested the normalization itself; Table 2), the reference workflows started from the same input file as OSMSS and normalized it in the same way (Figures 4A and 5A; Experimental Section). OSMSS and the reference workflows thus shared the input and differed in the statistical model and the analysis route. Because several graphical tools (Table 1) run the same statistical engines, such as limma, the reference workflows also stand in for these tools run with matched settings. We compared OSMSS with the reference workflows in two ways: by the range class, and by statistical tests against the median reference workflow, dataset by dataset. The range class says whether OSMSS’s value lies below, within or above the range of the reference workflows’ values. Above the range means more true positives, higher sensitivity, nMCC, pAUC or fold-change correlation, fewer false positives or a lower FDP than every reference workflow. Each benchmark has one main rule for calling a hit, either adjusted P alone or the fold-change filter, and reports the other rule as secondary (Table 2; Experimental Section). Among the protein benchmarks with several datasets, statistical tests that compared OSMSS with the median reference dataset by dataset found a significant difference, that is, an improved score, only in DDA. In DIA, TMT single plex and both TMT multi-plex designs, no metric was significant after adjustment under either rule (Experimental Section).

**Figure 4.**
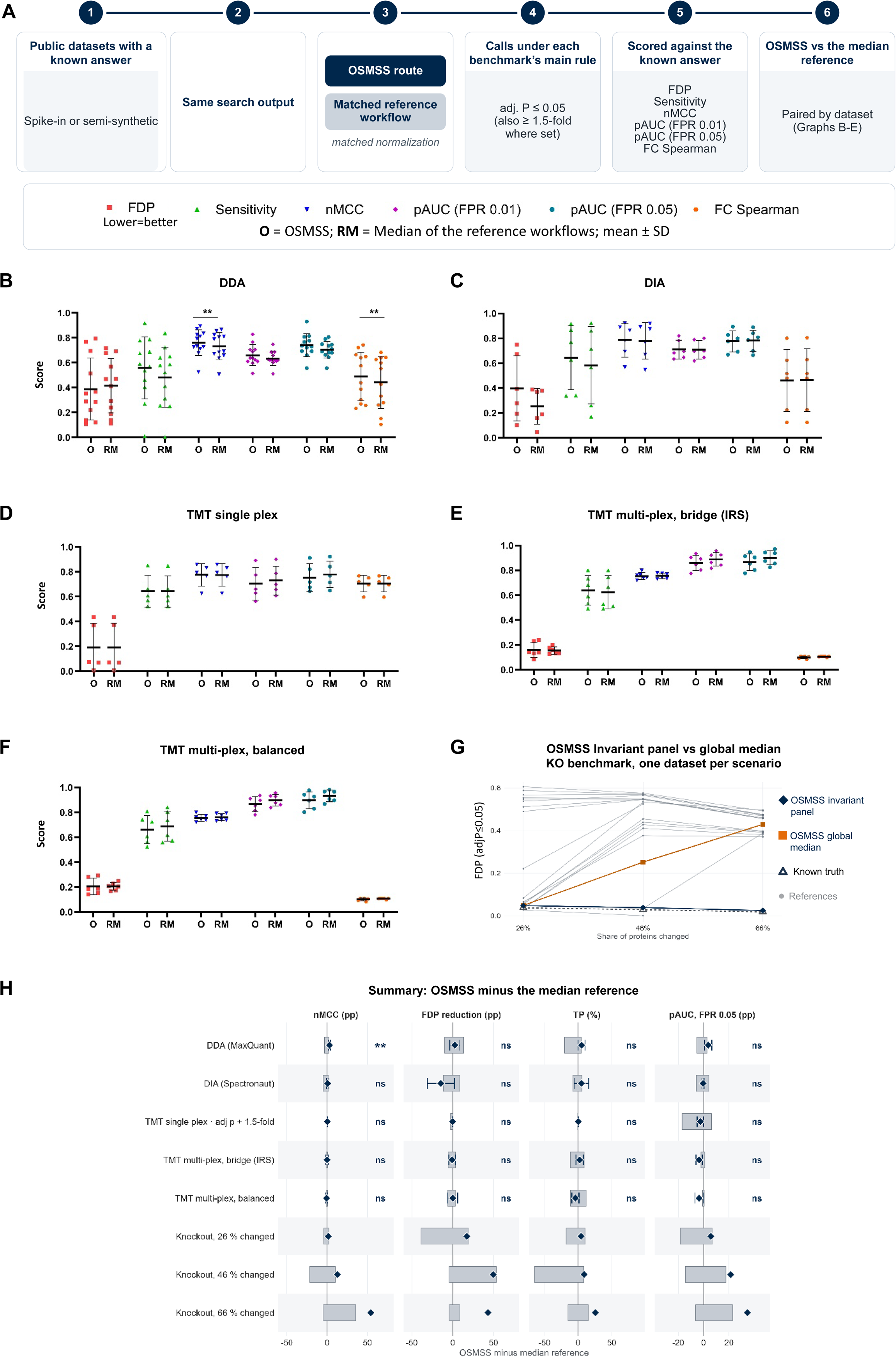
Global protein benchmarks: Comparison of OSMSS with matched reference workflows. (A) The benchmark design. (B–F) DDA, 12 datasets (B); DIA, 6 (C); TMT single plex, 5 (D); and TMT multi-plex, bridge (IRS) (E) and balanced (F) designs, 6 each. Calls are at adjusted P ≤ 0.05, plus ≥ 1.5-fold in D. O is OSMSS and RM the median reference; points are datasets, and lines and bars show mean ± SD. Six metrics are drawn on a 0 to 1 scale. The false-discovery proportion (FDP) is the share of calls that are false, and sensitivity the share of true changes called. The normalized Matthews correlation coefficient (nMCC) scores correct calls, wrong calls and non-calls together [117]. The partial area under the curve (pAUC), to false-positive rates (FPR) of 0.01 and 0.05, measures how well P values rank true changes first [118]. FC Spearman is the rank correlation of estimated with true fold changes. Stars mark two-tailed paired t-tests of OSMSS against the median reference. P values were Holm–Šidák-adjusted across eight metrics, including the undrawn true and false positive counts [119]. The thresholds are *P < 0.05, **P < 0.01, ***P < 0.001 and ****P < 0.0001 (adjusted); metrics without a star were not significant. (G) FDP at adjusted P ≤ 0.05 without the fold-change cut-off, by the share of proteins changed in the knockout simulation (invariant-protein normalization). Navy diamonds, OSMSS with the invariant-protein panel; orange squares, OSMSS with the global median; thin grey lines, reference workflows; dashed line with filled triangles (“Known truth”), normalization on the truly unchanged proteins, possible only with simulated data. (H) Summary: OSMSS minus the median reference. nMCC, FDP reduction and pAUC at the higher FPR are in percentage points, and true positives (TP) in % of the median reference. Positive FDP reduction means a lower FDP. Diamonds are means over datasets, bars unadjusted 95% confidence intervals and grey boxes the reference workflows’ range; “ns” (not significant) and stars are as in B–F.

**Figure 5.**
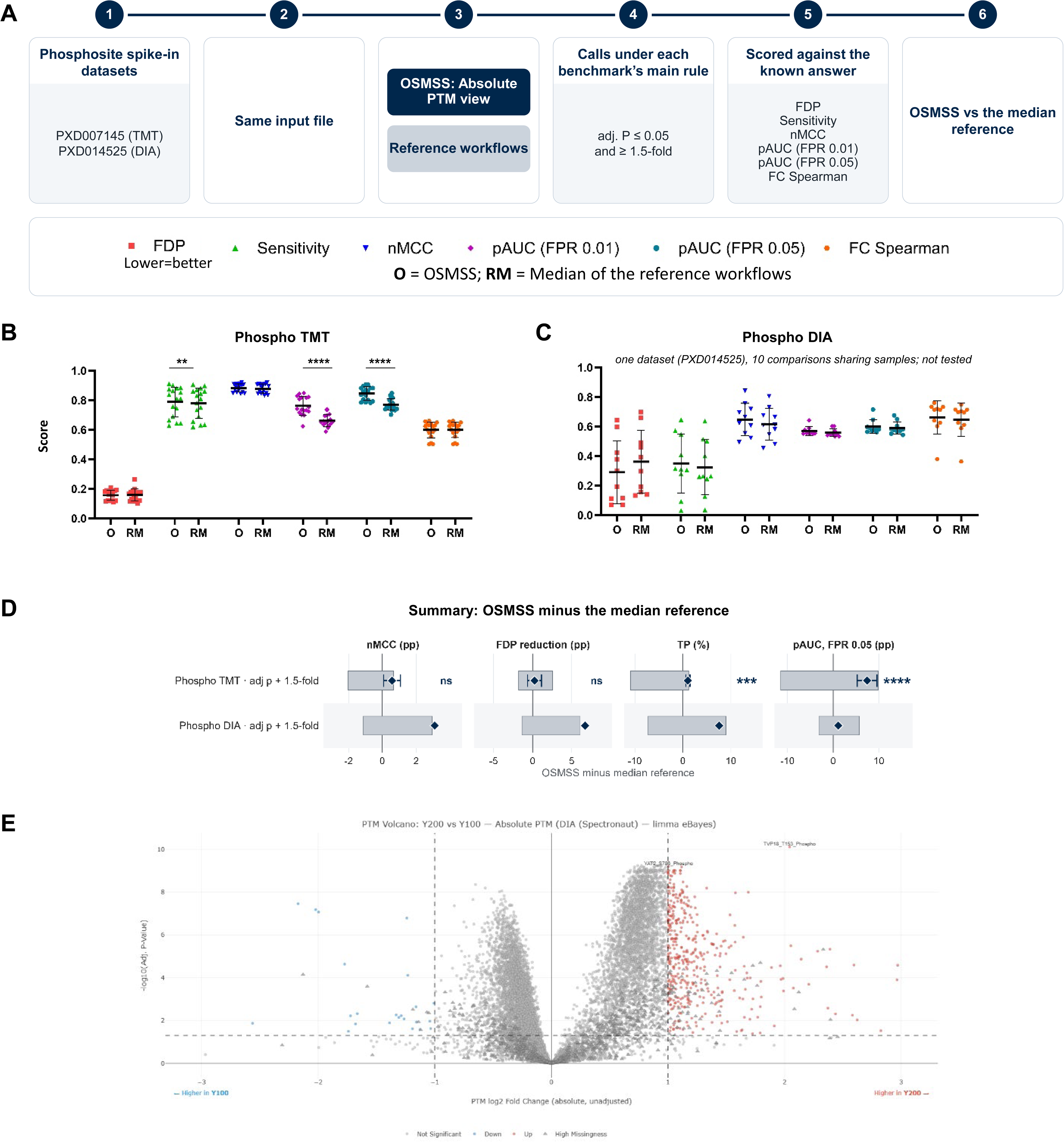
Phosphosite benchmarks: Comparison of OSMSS with matched reference workflows, in the Absolute PTM view (sites not adjusted for their parent protein). (A) The benchmark design, as in Figure 4A, on the phosphosite spike-in datasets PXD007145 (TMT) [111] and PXD014525 (DIA) [112]. Calls are at adjusted P ≤ 0.05 and ≥ 1.5-fold. (B) Phospho TMT, 17 datasets, one per TMT10 run, drawn as in Figure 4B–F. (C) Phospho DIA, one dataset with 10 comparisons that share samples. Points are comparisons, each paired with its own median of the reference workflows, and lines and bars show mean ± SD; this benchmark was not tested. (D) Summary: OSMSS minus the median reference, drawn as in Figure 4H. Stars for phospho TMT are from the paired t-tests in B. (E) Screenshot of a PTM volcano plot in OSMSS (Absolute view) for the phospho DIA dataset PXD014525 [112], Y200 vs Y100 (yeast phosphopeptides at twice the amount in a constant human background).

In label-free DDA, OSMSS identified more true changes than any reference workflow, while its false calls stayed within the reference range (Figure 4B). Against the median reference workflow, the differences were a few percentage points. In DIA, OSMSS fell within the reference range for true positives, sensitivity, nMCC and pAUC (Figure 4C). It was outside the range for false positives and FDP, with more false positives than every reference workflow. None of these differences from the median reference was significant after adjustment. The extra calls had small fold changes, and much of this excess comes from one setting in OSMSS’s limma model, the variance trend. In TMT single plex, with the fold-change filter as the main rule, OSMSS made as many true and false calls as the median reference workflow but ranked true changes slightly lower (Figure 4D). In TMT multi-plex, OSMSS was within the reference range on every call metric in both the bridge and the balanced designs, and it ranked true changes slightly lower than the references (Figure 4E,F). In the knockout benchmark, the invariant-protein panel kept the unchanged proteins centered at zero; global median normalization and the reference workflows moved them further off zero as the share of changed proteins grew. When 66% of proteins changed, global median normalization made 1,745 false calls, against 69 with the panel (Figure 4G shows the FDP in each scenario, Figure 4H summarizes the differences from the median reference).

In phospho TMT, with the fold-change filter as the main rule, OSMSS was within or above the reference range on every call metric (Figure 5B). Against the median reference, it found 1.1% more true changes and ranked them better. Without the filter, which is how OSMSS reports results by default, OSMSS found the most true changes but made 22.6% more false positives than the median reference. Most of these false positives were below 1.5-fold and in the direction in which median centering shifted the unchanged phosphosites. In phospho DIA, also with the fold-change filter as the main rule, OSMSS made the fewest false positives and had the lowest FDP of the seven workflows (Figure 5C). Without the filter, it made more false positives than the median reference, at almost the same FDP. Figure 5D summarizes the differences from the median reference for the two phosphosite benchmarks. Figure 5E shows a PTM volcano plot in OSMSS. Supporting Information Text gives the detailed results of each benchmark.

Two limits apply to this overview. The knockout and phospho DIA have one dataset per scenario, and therefore no interval or test. In the other protein benchmarks, with 5 to 12 datasets and eight metrics each, only large or consistent differences can reach significance after adjustment. In addition, the observed FDP at adjusted P ≤ 0.05 exceeded the nominal 5% for OSMSS and the reference workflows alike, a property of these spike-in designs (Supporting Information Text). Separately from these two limits, some parts of OSMSS could not be scored, because no public dataset with a known answer fits them. These are the Adjusted PTM view, for which datasets that can assess protein-corrected PTM analysis are scarce^82^, and multi-plex TMT PTM data. The Supporting Information Text describes these, and the other routes and inputs that were not benchmarked.

## Conclusions

OSMSS was designed so that bench scientists can access and apply established statistical models through questions about their experiment rather than through code. Users upload native exports from common proteomics software; their answers to OSMSS queries about the experimental design select an established route for normalization, missing values and the statistical model. Additional checks on the raw data, such as whether it has been normalized already, provide further guidance to the user on proceeding with analysis. Despite the increased user accessibility OSMSS provides through a graphical interface, the statistical performance it yields matched that of established reference workflows in our benchmarks while data handling and presentation are kept transparent to the user. Additional analyses provide information to the user regarding proteins that do not pass the filter, or do not fit well in the statistical test, such as proteins absent from one condition. Each analysis comes with a generated methods text, allowing users to better understand how their data were handled and to provide the text to a specialist when additional analysis is needed. The results connect to interpretation tools, from highlighting proteins on the output graphs, to gene set enrichment analysis and kinase activity inference with PhosR. Kinase prediction tools need no MS data and show both conservation and solvent accessibility alongside the AlphaFold structure, again favoring user ability to further explore their data. Analyses can be saved and easily shared among lab members and collaborators.

Benchmarking was conducted to assess whether the guidance provided by OSMSS compromises statistical robustness. Across seven benchmarks with known answers, OSMSS’s results were comparable to those of matched reference workflows, and no significant adverse changes were found in the benchmarks tested, under each benchmark’s main rule. Without a fold-change filter, OSMSS made more false positives than the median reference in phospho TMT, most of them small changes in the direction of the normalization shift. As such, OSMSS provides a guided, no-code user interface that allows analysis of proteomic search files with high transparency for users and a range of tools to explore and analyze private and publicly available proteomics data.

## Supporting Information

The Supporting Information is available alongside the publication: Figure S1, OSMSS code architecture; Supporting Information Text, detailed benchmark results, supplementary methods, and benchmark software and inputs (PDF); Supplemental Data 1, per-dataset benchmark values for OSMSS and every reference workflow (XLSX). The Spectronaut export schemes OSMSS_Global.rs and OSMSS_PTM.rs are provided in the OSMSS repository.

## Data and Software Availability

OSMSS is available as open-source software on GitHub (https://github.com/CoulombeLab/OSMSS; release v3.1, archived on Zenodo, doi:10.5281/zenodo.24170667), licensed under the GNU General Public License, version 3 or later. The repository includes guided installers for Windows, macOS and Linux. They check for R, install the R packages OSMSS needs, optionally set up SAINTexpress, The Kinase Library and PhosphoSitePlus from files the user downloads from their sources, and create a desktop shortcut that starts OSMSS. Because The Kinase Library matrices^48,49^ and the PhosphoSitePlus data^94^ have their own license terms, the repository does not include them. If they are not set up during installation, OSMSS directs the user to download them when a tool first needs them. The R packages OSMSS uses are free to use, are cited in the methods text generated by OSMSS, and are listed with links in the app’s external resources page. The web services OSMSS calls (14-3-3-Pred^96^ and Scansite^95^) are credited in the graphs they produce. OSMSS also queries UniProt, the AlphaFold Protein Structure Database and the Ensembl REST service; only identifiers or protein sequences are sent to these services, never quantitative data. No new mass spectrometry data were generated for this study. All benchmarks reused public data: the OpDEA spike-in datasets^13^, the MSstatsTMT spike-in mixtures^80^, timsTOF Pro HeLa runs from Kalxdorf et al.^10^, and the phosphoproteomics datasets PXD007145^111^ and PXD014525^112^. The Benchmark Datasets subsection of the Experimental Section gives the source of each dataset.

## Acknowledgments

We thank Dr. Venkatesha Basrur and Dr. Alexey Nesvizhskii, and the staff of the Proteomics Resource Facility at the University of Michigan, for their long-standing collaboration and advice, and for the analyses and search files from earlier experiments that helped shape OSMSS. We thank the laboratories of Drs. Carole Parent, Rachel Zemans, Diane Fingar, Ben Allen, Chelsey Spriggs, Hisham Bazzi and Kristen Verhey, as well as members of the Coulombe laboratory, for testing OSMSS on their own datasets and for their feedback. We also thank Craig N. Johnson for his assistance with IT services and for his advice.

## AI Disclosure

AI tools (Claude Code Opus and Sonnet models, 4.8, 5.0 and 5.5; GitHub Copilot) assisted in developing OSMSS’s code alongside manual code using VS Code. At every stage of development, a human (author EC) audited the code. OSMSS itself calls no language model, so neither users nor their data interact with AI models. Its methods text is generated by fixed rules. The authors used Claude models (Anthropic, Opus and Sonnet 5.5 models) during the preparation of this manuscript to synthesize initial drafts, improve language, correct grammatical errors, refine prose and perform review of citations used and statements. After using these tools, the authors reviewed, edited, and revised the content to ensure accuracy of language.

## Author Contributions

E.C.: Conceptualization, Methodology (lead), Software, Validation, Formal analysis, Investigation, Data curation, Visualization, Writing – original draft, Writing – review & editing, Project administration. P.A.C.: Conceptualization, Methodology (supporting), Supervision, Resources, Funding acquisition, Writing – review & editing. All authors have given approval to the final version of the manuscript.

