## Supplemental Material and Figure S1 for "One Stop MS Shop (OSMSS): statistical analysis and interpretation of quantitative proteomics data for non-coders"

This file contains detailed benchmark results (label-free DDA; DIA and TMT; phosphosites; limitations), supplementary methods for the analysis routes and the interpretation tools, and the benchmark software and inputs, and Figure S1 (OSMSS code architecture). Table1, Table2, and Supplemental Data 1 (per-dataset benchmark values for OSMSS and every reference workflow, XLSX) are provided as separate files.

### Detailed Benchmark Results

#### Label-free DDA

In label-free DDA, OSMSS found more true changes than any reference workflow, while its false calls stayed within the reference range (Figure 4B). The benchmark used 12 spike-in datasets (22 comparisons), read from MaxQuant raw Intensity. OSMSS was above the reference range on 6 of 8 metrics, among them true positives, sensitivity, nMCC and pAUC(0.05). Against the median reference workflow, the differences were a few percentage points. OSMSS's nMCC was 2.8 pp higher (95% CI +1.4 to +4.2; adjusted P = 0.0089). Its fold-change correlation was 4.9 pp higher (+2.4 to +7.4; adjusted P = 0.0089).

Looking at each dataset separately, OSMSS was within or above the reference range in 93 of 96 per-dataset results. The three below the range were in two dilution series of UPS1[1] in a yeast background[2,3], in which OSMSS utilizing proDA fitted 5 and 9 conditions in one model. In these series, OSMSS made more false positives than the references (73 vs at most 46.5 and 103 vs at most 77), but also identified the most true changes (31.5 vs at most 16.5, and 22.7 vs at most 15.3). We attribute this to fitting all conditions in one model, which gives the test more residual degrees of freedom, that is, more data from which to estimate variability. Despite OSMSS scoring within or above the reference range in 96.9% of per-dataset results, no difference in true positives, false positives or FDP was significant after adjustment.

The knockout benchmark tested the invariant-protein panel where global normalization is expected to fail: when many proteins change in one direction. In its semi-synthetic data (Experimental Section), rescaling each knockout run to the control runs' total protein moved the unchanged proteins off zero, and more so as the share of changed proteins grew. Without normalization, their median log2 fold change was +0.045, +0.255 and +0.539 at 26%, 46% and 66% changed. OSMSS's panel held only truly unchanged proteins in every scenario. With no change (0%), OSMSS made 8 false calls, 7 of them flagged as LowEvidence, against 0 to 46 for the reference workflows. With the panel, the unchanged proteins stayed centered at zero, with a median log2 fold change of −0.006 to −0.008. Global median normalization left them at +0.727 at 66%, and the reference workflows at +0.537 to +0.964. At 26% changed, OSMSS was within the reference range on every metric. As the share of changed proteins grew, OSMSS's nMCC exceeded the median reference's by 12.7 pp at 46% and by 54.4 pp at 66%. The difference from its own global median normalization widened too. At 26% changed, the two gave nearly the same result (50 and 51 false calls; false calls are false positives plus calls in the wrong direction, Supplemental Data 1), and at 46% global median normalization made 628 false calls against 75 with the panel. At 66%, global median normalization made 1,745 false calls, against 69 with the panel. The corresponding FDPs were 0.429 and 0.025 (Figure 4G). At 66%, OSMSS's sensitivity was 0.916. These measurements describe the panel's behavior under a large one-directional shift and affirm its use in specific experimental contexts where global median normalization is assumed to fail.

**DIA and TMT**

In DIA, OSMSS fell within the reference range for true positives, sensitivity, nMCC and pAUC, and outside it for false positives and FDP (Figure 4C). None of these differences was significant. Against the most liberal reference, limma, OSMSS made 9.2% more false positives (adjusted P = 0.14), and its FDP was 3.8 pp higher (adjusted P = 0.19). Against the median reference workflow, it made 19.5% more false positives (adjusted P = 0.32), and its FDP was 14.4 pp higher (adjusted P = 0.37). The extra calls had small effects. In the three datasets we examined in detail (HEof_n600, HEof_w600 and HYtims134), their median absolute log2 fold change was 0.17–0.29, or about 1.1- to 1.2-fold. Much of this excess comes from one setting in OSMSS's limma model, the variance trend. OSMSS runs limma's empirical Bayes test with the trend, which lets the expected variance change with protein intensity. The limma reference workflow runs the same test without it. When we turned the trend off in OSMSS (Experimental Section, DIA route), false positives across the six DIA datasets fell by 15% and true positives fell by 2.3%. We kept the trend despite this. In proteomics data, low-abundance proteins are usually measured less precisely than abundant ones. The trend models this[4], and the Bioconductor expression-proteomics workflow uses it[5]. We chose the setting suited to typical experimental data rather than the one that scores best on these benchmarks. The gap narrowed with a 1.5-fold cutoff (|log2 FC| > 0.58), a threshold many users apply when reading volcano plots. With this cutoff, OSMSS made 11.5% more false positives than the median reference (adjusted P = 0.34), and its FDP was 9.1 pp higher (adjusted P = 0.69). Its FDP stayed just outside the reference range (0.204 vs 0.089–0.185) and was not significantly higher than that of limma, the reference at the top of the range (adjusted P = 0.85).

In TMT single plex, OSMSS made as many true and false calls as the median reference workflow but ranked true changes slightly lower. This benchmark (five datasets, 15 comparisons) is scored with the fold-change filter. Here, median centering shifted the unchanged human background by a median log2 fold change of −0.267, so without the filter every workflow's FDP was between 0.55 and 0.71. With the filter, OSMSS was within the reference range on 7 of 8 metrics and above it on nMCC by a margin of about 0.001 (Figure 4D). Dataset by dataset, 38 of 40 per-dataset results were within or above the range; the 2 below were fold-change correlations short by about $10^{-6}$. Against the median reference, true positives differed by +0.2%, false positives by 0.0% and FDP by 0.0 pp. The pAUC(0.05) was 2.5 pp lower (95% CI −5.1 to −0.03; adjusted P = 0.29), and no metric was significant after adjustment. Here as well, the lower pAUC traces to the variance trend and to fitting all conditions together, both recommended parameters, and neither changes a call.

Without the trend, OSMSS's pAUC(0.05) was 0.789 rather than 0.754. Fitting only the two compared conditions gave 0.785. True positives and FDP with the fold-change filter stayed the same in every variant. Without the fold-change filter, OSMSS made 15.2% more false positives than the median reference workflow (95% CI +2.0 to +28.4; adjusted P = 0.21), not significant after adjustment.

In TMT multi-plex (six five-plex datasets, 36 comparisons), OSMSS was within the reference range on every call metric in both the bridge and the balanced designs (Figure 4E,F). It ranked true changes slightly lower than the references. Its pAUC(0.05) was 0.866 against 0.880–0.908 in the bridge design, and 0.896 against 0.924–0.937 in the balanced design. No difference from the median reference was significant after adjustment. OSMSS's pAUC(0.05) was 3.5 and 3.6 pp lower than the median reference in the two designs (adjusted P = 0.12 and 0.23), and its nMCC was 0.1 and 0.6 pp lower. It made about 21% more false positives in both designs (adjusted P = 0.35 and 0.46). In absolute terms, this is less than one extra false positive per comparison: 2.8 against 2.3 in the bridge design, and 4.1 against 3.4 in the balanced design.

The slightly lower ranking has two sources. First, OSMSS tests a protein only if it is quantified in at least half of the plexes (the coverage rule described in the Experimental Section, TMT route). This left at most one UPS1 protein untested per MS2 dataset, and 1, 7 and 5 in the three SPS-MS3 datasets, all listed in OSMSS's not-tested table provided to the user. The coverage rule accounted for about 0.02–0.03 of the lower pAUC(0.05). On the proteins that every workflow tested, OSMSS's pAUC(0.05) was within the range (0.916 against 0.915–0.939, bridge design). Second, fitting all conditions in one model lowered pAUC(0.05) by about 0.02. Ratio compression was similar for OSMSS and the references.

**Phosphosites**

For phospho TMT we used yeast phosphopeptides diluted into a constant HeLa background[6] (Figure 5A). The 17 TMT10 runs gave 51 comparisons, scored in the Absolute view. About 17% of sites rise in this design, and median centering therefore shifts the unchanged human sites by 0.16–0.35 log2. TMT replicates are precise enough to detect this small shift. With an adjusted P alone, every workflow's FDP was high (0.52–0.72 for the references). We therefore used a 1.5-fold filter as the main rule, as for TMT single plex. With the filter, OSMSS was within or above the reference range on every call metric (Figure 5B) and in 124 of 136 per-dataset results. Against the median reference, it found 1.1% more true changes (adjusted P = 0.0006) and ranked them better (pAUC(0.05) 7.4 pp higher; adjusted P = $1.3 \times 10^{-5}$). It made 87 false positives per comparison against 89 for the median reference, and neither false positives nor FDP differed (adjusted P = 0.76 for both).

Without the filter, which is how OSMSS reports results by default, OSMSS found the most true changes but made 22.6% more false positives than the median reference (adjusted P = $1.0 \times 10^{-5}$). Its FDP was 0.73 against 0.52–0.72 for the references (6.7 pp above the median; adjusted P = $1.0 \times 10^{-6}$). Of these false positives, 94% were below 1.5-fold and 94% were in the direction of the centering shift, as were 86–94% and 87–94% of the references'. OSMSS was close to the limma reference (1,327 against 1,300 false positives per comparison).

In phospho DIA, the benchmark uses PXD014525[7], one dataset of 10 comparisons that share samples. Here 37% of sites rise, and Spectronaut's cross-run normalization shifted the unchanged human sites by −0.27 to −0.31 log2 on average. With an adjusted P alone, every workflow's FDP was therefore high (0.49–0.61), and thus we applied the same 1.5-fold filter as for phospho TMT.

With the filter, OSMSS made the fewest false positives of the seven workflows (1,000 per comparison against 1,052–1,286 for the references) and had the lowest FDP (0.291 against 0.297–0.371; Figure 5C). Against the median reference, it made 16.0% fewer false positives and found 7.6% more true changes. OSMSS was within or above the reference range in 77 of 80 per-comparison results. The three below the range were all in the comparison whose true 1.5-fold change sits exactly at the filter threshold. OSMSS combines each site's precursors by median polish, whereas the reference workflows summed them; median polish compresses ratios slightly more: its measured yeast fold changes reached 77% of the expected values, against 84% for the references that summed precursors and 78% for MSstatsPTM. Without the filter, OSMSS found more true changes than any reference (2,492 per comparison against 1,818–2,470). It made 16.1% more false positives than the median reference and 12% more true positives, and its FDP was therefore almost the same (0.603 against 0.600).

**Limitations: where no benchmark with a known answer exists**

Some parts of OSMSS could not be scored, because no public dataset with a known answer fits them. The first is the Adjusted PTM view. A change in a phosphosite combines changes in the protein and in its phosphorylation[8-10]. The published adjustments, and OSMSS's own, are described in the main text (Results and Discussion) and the Experimental Section (PTM route). Datasets that can assess protein-corrected PTM analysis are scarce[10]. The one we found, the MSstatsPTM ubiquitination spike-in[9], is label-free DDA, which OSMSS's PTM routes do not read. In the two phosphosite dilution series, the protein-level (global) files that the Adjusted view needs are stand-ins built from the enriched data, so scoring the Adjusted view against them would be circular. The second is multi-plex TMT PTM data. To our knowledge, no public phosphosite dataset with a known answer spans several plexes. OSMSS's multi-plex PTM route offers "Multi-Plex + IRS Bridge Normalization" and "Multi-Plex (Plex as Run Covariate)", the two designs benchmarked at the protein level as bridge and balanced. Last, phospho DIA rests on one dataset, because PXD014525 is, to our knowledge, the one public DIA phospho dilution series with a known answer[7]. Other parts of OSMSS were not benchmarked separately, because they run an established tool unchanged or feed the benchmarked models from other inputs. These are the affinity-purification route, which runs SAINTexpress as published; label-free DDA input from Proteome Discoverer and FragPipe (the DDA benchmark used MaxQuant output); DIA-NN input (the DIA benchmark used Spectronaut output); the N=1 module; the interpretation tools; and the invariant-protein panel on real knockout data.

Across the benchmarks, the observed FDP at adjusted P ≤ 0.05 was well above the nominal 5%, for OSMSS and the reference workflows alike (for example, 0.39 for OSMSS against 0.35–0.45 for the references in DDA). This is a property of the spike-in designs rather than of any one workflow. Mixing and loading are never exact, normalization can shift the unchanged background, shared peptides can carry signal across species, and with several replicates per condition even small systematic shifts reach significance, whereas the Benjamini–Hochberg procedure controls the false discovery rate only when unchanged features are truly unchanged[11]. For users, this is a reason to read adjusted P values together with fold changes, as the OSMSS volcano plot does by coloring proteins above a fold-change threshold.

### Supplementary Methods: analysis routes

The invariant-protein panel is chosen from proteins detected in every run, including control runs. A protein qualifies when its abundance is at or above the 75th percentile, its standard deviation within each condition is at most 0.15 log2, and its difference between conditions is at most 0.1 log2 once the differences are re-centered on their mode (the offset shared by the bulk of proteins) rather than on zero. If too few proteins qualify, OSMSS relaxes these limits stepwise, up to a standard deviation of 0.30 log2 and a difference of 0.5 log2, and then to the top 50% of proteins by abundance. A panel of 10 to 19 proteins is used with a low-confidence warning; with fewer than 10, OSMSS falls back to global median normalization. Panel proteins stay in the model and are tested; because they were selected as stable, their fold changes are close to zero by design. Besides the knockout answer described in the main text, the wizard selects the panel for affinity-purification controls from another cellular fraction or the same bait in another condition, and for a non-specific capture control in a factorial design, which is analyzed with proDA. In the knockout benchmark, the panel held 986, 749, 575 and 369 proteins at 0%, 26%, 46% and 66% changed, and all were truly unchanged. These scenarios are stress tests of a large one-directional shift rather than models of a single knockout and, being semi-synthetic, they test the panel as a proof of concept.

For MaxQuant input, contaminant protein groups are kept by default and can be removed in the wizard. For Proteome Discoverer input, OSMSS reads the raw Abundance columns and ignores Proteome Discoverer's normalized and scaled abundances. For FragPipe input with IonQuant normalization on, OSMSS does not apply the global median, but applies the invariant-protein panel when the wizard selects it. When the user assigns batches or paired samples, the batch or the subject enters the proDA and limma models as a covariate (in limma, paired samples are handled with duplicateCorrelation[4]). Batch correction in the PCA and heatmap views (removeBatchEffect in limma) changes only the display, not the tested values.

OSMSS does not normalize intensities before SAINTexpress. When the largest intensity exceeds $10^8$, all intensities are divided by one common factor so that the largest is $10^8$, and detected values below 1 are set to 1, which keeps the ratios between purifications. A prey needs at least two values, and preys matching the bait gene are removed. Interactors are called at a SAINTexpress Bayesian false discovery rate of at most 0.05, and SaintScores of at least 0.8 and 0.5 are labeled high and medium confidence. When the bait or the control has fewer than two replicates, or the design is factorial, OSMSS uses proDA instead.

In the DIA protein route, before the detection filter described in the main text, OSMSS removes proteins quantified in fewer than 50% of the samples of every condition. These proteins are neither fitted nor tested for presence or absence, and their number is given in the methods text.

The run-median check calls a spread between 0.02 and 0.03 log2 borderline, and it is inconclusive when fewer than 100 proteins (or 5% of all proteins, if more) are quantified in every run; in every case the check is advisory, and the checkbox decides whether OSMSS normalizes. k-nearest-neighbor imputation uses k = 10 and fills only proteins missing at most one value in each of the two compared conditions. For the choice between one model and a two-condition fit, the typical within-condition standard deviation of each condition with at least three samples is the median of its proteins' standard deviations; when fewer than two conditions can be assessed, only the two compared conditions are fitted. In the NMAR tab, fold changes are estimated after imputation from a down-shifted normal distribution (1.8 standard deviations below the mean, width 0.3 standard deviations), for display and ranking only. The Fisher test itself uses detection counts, and returns no P value when even complete separation of the two conditions could not reach $P \leq 0.05$.

Both PTM routes currently analyze phosphosites. The Protein view fits limma to the parent proteins of the global file, with the variance trend and robust estimation. Phosphosites that fail the detection filter are tested by Fisher's exact test on detection counts in the Absolute view, and receive no P value when they cannot reach $P \leq 0.05$.

### Supplementary Methods: interpretation tools

Trend clusters group the condition means of each protein, scaled as z-scores, by fuzzy c-means clustering (e1071)[12]. The fuzzifier is estimated from the numbers of proteins and conditions by the method of Schwämmle and Jensen[13], as in Mfuzz[14]. Kinase-Site Analysis is OSMSS's implementation of PhosR's kinase–substrate scoring[15], with PhosR's kinase motifs and substrate sets and, for TMT data, RUVphospho correction; it reads the PhosR-format file exported from PTM DE. Kinase-substrate enrichment analysis itself is not run within OSMSS; PTM DE exports a file formatted for it[16,17]. Gene set enrichment analysis uses gene sets of 15 to 500 genes and at least 10,000 permutations, on human or mouse MSigDB gene sets matched by gene symbol, downloaded through msigdbr[18] (version 26.1.1 here) when first used and cached with their download date, and ranks genes by the user's .rnk file, by a limma statistic computed from .gct and .cls files, or by the π-score of the OSMSS differential expression export. Solvent accessibility is computed by OSMSS's own implementation of the Shrake–Rupley method[19] (300 points per atom and a 1.4 Å probe; absolute accessible area in $Å^2$) on the AlphaFold model. The model's per-residue confidence (pLDDT) is not used, so accessibility in low-confidence regions, which are often disordered, should be read with care. Conservation is the fraction of orthologs whose aligned residue matches the human residue, across 11 species: gorilla, marmot, leopard, polar bear, pig, horse, dolphin, blue whale, opossum, turkey and tortoise. The PTM database stores sites as reported in FragPipe's psm.tsv and peptide.tsv files, after FragPipe's own false discovery rate filtering, and applies no further filter. Saved Experiments hold R data (RDS) files; users should open only files from trusted sources and use R 4.4 or later, which corrects a vulnerability in loading R data files.

### Supplementary Methods: benchmark software and inputs

OSMSS benchmark results were produced with OSMSS's own code under R 4.1.2 and limma 3.50.1 on our server, or on a local computer without the interface, with limma's legacy empirical Bayes setting (eBayes(legacy = TRUE)) to match the server; the two reproduced each other (largest difference in adjusted P below $10^{-12}$, no discordant calls). Reference workflows ran under R 4.6.1, with ROTS 2.4.0, DEqMS 1.30.0, DEP 1.32.0, samr 3.0.1, MSstatsTMT 2.20.0 and MSstatsPTM 2.14.0. The OpDEA datasets were used as searched by OpDEA[20], with MaxQuant 2.1.0.0 (DDA), MaxQuant 2.4.4.0 (TMT single plex) and Spectronaut 18.3 (DIA); the TMT multi-plex PSMs were from Proteome Discoverer 2.2[21]. OSMSS has no MaxQuant TMT reader. For TMT single plex, each dataset's MaxQuant proteinGroups.txt and evidence.txt were therefore converted to Proteome Discoverer protein and nested peptide files, whose abundances equaled OpDEA's in every shared value. For phospho TMT, the authors' MaxQuant 1.5.5.4 modificationSpecificPeptides.txt was converted to a Proteome Discoverer phosphopeptide-group file, with MaxQuant's corrected reporter intensities and each site's localization taken from its best-scoring PSM; every workflow received the same file. Neither phosphosite study measured an unenriched proteome, so the global file the wizard requires was a stand-in: for phospho TMT, the sum of each run's peptides per protein, and for phospho DIA, Spectronaut's protein-group report from the same enriched runs. The stand-ins enter only the Adjusted view, which was not scored. In phospho DIA, every workflow used the same Spectronaut 21.1 precursor report; the reference workflows summed precursor intensities per site, and MSstatsPTM used the precursors directly. In TMT multi-plex (MassIVE.quant RMSV000000266 for MS2 and RMSV000000265 for SPS-MS3), every workflow received one peptide table, produced once from the Proteome Discoverer PSMs by MSstatsTMT's converter. The knockout benchmark used the six PXD021425 runs with 4.5% and 6% E. coli spike-ins (three each; the authors' MaxQuant 1.6.14 search), from which all E. coli protein groups were removed, so that only the HeLa background remained, and the three HYpro16 acquisitions are from PXD020815.

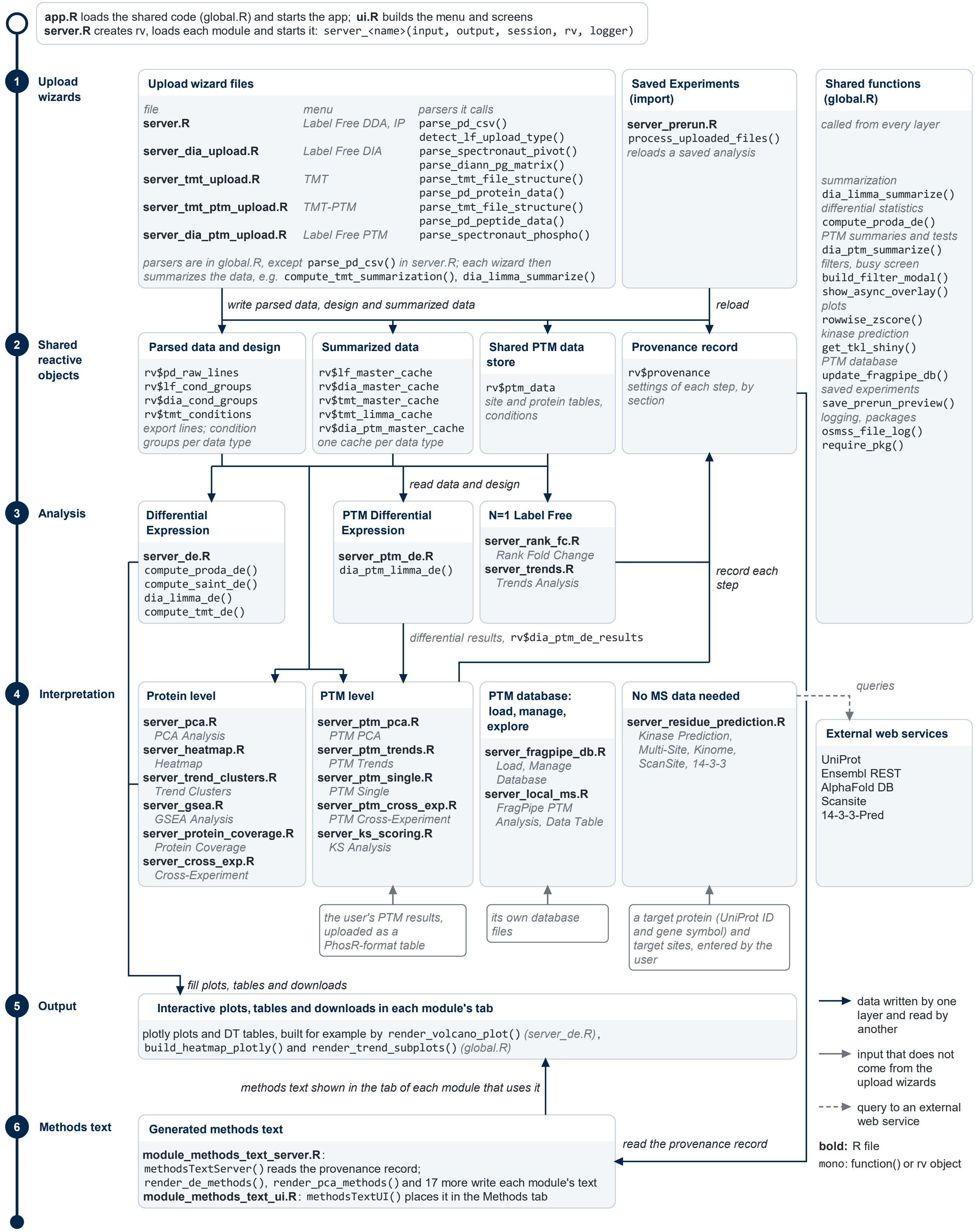

app.R loads the shared code (global.R) and starts the app; ui.R builds the menu and screens
server.R creates rv, loads each module and starts it: server_<name>(input, output, session, rv, logger)
1 Upload wizards
Upload wizard files
file
menu
parsers it calls
server.R
Label Free DDA, IP
parse_pd_csv()
detect_lf_upload_type()
server_dia_upload.R
Label Free DIA
parse_spectronaut_pivot()
parse_diann_pg_matrix()
server_tmt_upload.R
TMT
parse_tmt_file_structure()
parse_pd_protein_data()
server_tmt_ptm_upload.R
TMT-PTM
parse_tmt_file_structure()
parse_pd_peptide_data()
server_dia_ptm_upload.R
Label Free PTM
parse_spectronaut_phospho()
parsers are in global.R, except parse_pd_csv() in server.R; each wizard then summarizes the data, e.g. compute_tmt_summarization(), dia_limma_summarize()
Saved Experiments (import)
server_prerun.R
process_uploaded_files()
reloads a saved analysis
Shared functions (global.R)
called from every layer
summarization
dia_limma_summarize()
differential statistics
compute_proda_de()
PTM summaries and tests
dia_ptm_summarize()
filters, busy screen
build_filter_modal()
show_async_overlay()
plots
rowwise_zscore()
kinase prediction
get_tkl_shiny()
PTM database
update_fragpipe_db()
saved experiments
save_prerun_preview()
logging, packages
osmss_file_log()
require_pkg()
write parsed data, design and summarized data
reload
2 Shared reactive objects
Parsed data and design
rv$pd_raw_lines
rv$lf_cond_groups
rv$dia_cond_groups
rv$tmt_conditions
export lines; condition groups per data type
Summarized data
rv$lf_master_cache
rv$dia_master_cache
rv$tmt_master_cache
rv$tmt_limma_cache
rv$dia_ptm_master_cache
one cache per data type
Shared PTM data store
rv$ptm_data
site and protein tables, conditions
Provenance record
rv$provenance
settings of each step, by section
read data and design
3 Analysis
Differential Expression
server_de.R
compute_proda_de()
compute_saint_de()
dia_limma_de()
compute_tmt_de()
PTM Differential Expression
server_ptm_de.R
dia_ptm_limma_de()
N=1 Label Free
server_rank_fc.R
Rank Fold Change
server_trends.R
Trends Analysis
record each step
differential results, rv$dia_ptm_de_results
4 Interpretation
Protein level
server_pca.R
PCA Analysis
server_heatmap.R
Heatmap
server_trend_clusters.R
Trend Clusters
server_gsea.R
GSEA Analysis
server_protein_coverage.R
Protein Coverage
server_cross_exp.R
Cross-Experiment
PTM level
server_ptm_pca.R
PTM PCA
server_ptm_trends.R
PTM Trends
server_ptm_single.R
PTM Single
server_ptm_cross_exp.R
PTM Cross-Experiment
server_ks_scoring.R
KS Analysis
PTM database: load, manage, explore
server_fragpipe_db.R
Load, Manage Database
server_local_ms.R
FragPipe PTM Analysis, Data Table
No MS data needed
server_residue_prediction.R
Kinase Prediction, Multi-Site, Kinome, ScanSite, 14-3-3
queries
External web services
UniProt
Ensembl REST
AlphaFold DB
Scansite
14-3-3-Pred
the user's PTM results, uploaded as a PhosR-format table
its own database files
a target protein (UniProt ID and gene symbol) and target sites, entered by the user
fill plots, tables and downloads
5 Output
Interactive plots, tables and downloads in each module's tab
plotly plots and DT tables, built for example by render_volcano_plot() (server_de.R), build_heatmap_plotly() and render_trend_subplots() (global.R)
data written by one layer and read by another
input that does not come from the upload wizards
query to an external web service
bold: R file
mono: function() or rv object
methods text shown in the tab of each module that uses it
6 Methods text
Generated methods text
module_methods_text_server.R:
methodsTextServer() reads the provenance record;
render_de_methods(), render_pca_methods() and 17 more write each module's text
module_methods_text_ui.R: methodsTextUI() places it in the Methods tab
read the provenance record

**Figure S1.** Code layers of OSMSS. The left ribbon numbers the six layers of the code. In each layer, cards name the R files that make it up (bold) and, in monospace, their main functions or the shared reactive objects (rv) they write and read: (1) the upload wizards, with their menu items and the parsers they call; (2) the rv objects created in server.R; (3) the model functions of the analysis modules; (4) the interpretation modules, with their menu items; (5) example plot builders; and (6) the methods-text module. Solid arrows, data written by one layer and read by another; grey arrows, input that does not come from the upload wizards; dashed arrow represent queries to external web services. The side column lists the purpose groups of the shared functions in global.R, with examples.
